# Supplemental vibrotactile feedback of real-time limb position enhances precision of goal-directed reaching

**DOI:** 10.1101/327049

**Authors:** Nicoletta Risi, Valay Shah, Leigh A. Mrotek, Maura Casadio, Robert A. Scheidt

## Abstract

We examined vibrotactile stimulation as a form of supplemental limb state feedback to enhance on-going control goal-directed movements. Subjects wore a two-dimensional vibrotactile display on their non-dominant arm while performing horizontal planar reaching movements with their dominant arm. The vibrotactile display provided feedback of hand position such that small hand displacements were more easily discriminable using vibrotactile feedback than with intrinsic proprioceptive feedback. When subjects relied solely on proprioceptive feedback to capture visuospatial targets, target capture performance was degraded by proprioceptive drift and an expansion of task space. By contrast, reach accuracy was enhanced immediately when subjects were provided vibrotactile feedback, and further improved over two days of training. Improvements reflected a resolution of proprioceptive drift which occurred only when vibrotactile feedback was active, demonstrating that the benefits of vibrotactile feedback are due in part to its integration into the ongoing control of movement. A partial resolution of task space expansion that persisted even when the vibrotactile feedback was inactive demonstrated that training with vibrotactile feedback also induced changes in movement planning. However, the benefits of vibrotactile feedback come at a cognitive cost. All subjects adopted a stereotyped, movement decomposition strategy wherein they attempted to capture targets by moving first along one axis of the vibrotactile display and then the other. For most subjects, this inefficient movement approach did not resolve over two bouts of training performed on separate days, suggesting that additional training is needed to fully integrate vibrotactile feedback into the planning and online control of goal-directed reaching.

## Introduction

Information arising from limb proprioceptors contributes importantly to the planning and ongoing control of movements (Sainburg et al., 1993; Sanes et al., 1984; Scheidt et al., 2005; Sober & Sabes, 2003). Proprioceptors, predominantly muscle spindle afferents (Gandevia, et al., 1992; Matthews, 1988; c.f. Proske & Gandevia, 2012), give rise to sensations of limb position and movement. This is termed kinesthesia (Bastian, 1887), which is essential to carry out activities of daily life. Unfortunately, many people – including almost 50% of stroke survivors – experience deficits of kinesthesia in at least one arm (Carey and Matyas, 2011; Connell et al., 2008; Dukelow et al., 2010). These sensation deficits contribute to impaired control of reaching and stabilization behaviors that are vital to an independent life style (Blennerhassett et al., 2007; Scheidt and Stoeckmann, 2007; Tyson et al., 2008; Zackowski et al., 2004). Although such individuals often rely on visual feedback to guide movement, lengthy delays of visual processing (100-200 ms; (Cameron et al., 2014) yield slow, poorly-coordinated actions that require focused attention (Ghez et al., 1995; Sainburg et al., 1993); visually guided corrections come too late and result in jerky, unstable movements (Sarlegna et al., 2006).

Several research groups have proposed technological solutions to problems caused by somatosensory deficits, with notable examples being robotic re-training of proprioception (Cuppone et al., 2015; Cuppone et al., 2016; De Santis et al., 2015; Wong et al., 2011) and the application of supplemental performance feedback using vibrotactile stimulation (c.f., Bark et al., 2011; Krueger et al., 2017; Lieberman and Breazeal, 2007; Tzorakoleftherakis et al., 2015). Building on that foundation, a long-term goal of our work is to re-establish or enhance closed-loop control of goal-directed behaviors in individuals with impaired kinesthesia by creating sensory substitution technologies that provide real-time feedback of the moving limb’s state (e.g., position and velocity of the arm and hand) to a site on the body retaining somatosensation (such as the ipsilesional arm).

We also consider the possibility of enhancing skilled manual performance in a variety of applications including teleoperation tasks such as robotic surgery, where visual attention is constrained and precision of manual movements is desired. Previously, Krueger and colleagues showed that neurologically intact control subjects were able to enhance the accuracy and precision of goal-directed stabilization and reaching tasks performed with their dominant arm after only 3 to 5 minutes of practice with a vibrotactile sensory substitution system that applied feedback of dominant arm motion to the non-moving arm and hand (Krueger et al., 2017). A first set of experiments found that a limb state feedback scheme that encoded predominantly hand position information minimized performance errors during stabilization and reaching tasks. A second set of experiments compared this optimal limb state feedback to hand position error feedback – one of the simplest forms of “goal aware” feedback (Tzorakoleftherakis et al., 2015) – to determine the performance benefits of each encoding scheme. Both forms of synthesized feedback were able to enhance performance of stabilization and reaching behaviors in the absence of visual feedback. Although error encoding yielded superior results, likely due to the inclusion of information related to the spatial goal of movement, there are technological challenges associated with estimating the user’s motor intentions in non-structured, dynamically changing environments. Therefore, based on the finding that subjects can use limb state feedback to enhance the control of stabilizing and reaching actions throughout the arm’s workspace, we sought to advance our long-term goal by understanding the sensorimotor learning that leads to improved control of arm movements guided by supplemental limb state feedback systems.

Here, we characterized how the human sensorimotor system may use supplemental limb state feedback to enhance the on-going control of a moving limb. In particular, we assessed the extent to which supplemental vibrotactile feedback can be used to enhance the control of arm movements beyond performance limits imposed by intrinsic proprioceptive feedback. We exploited the fact that the vast majority of people – including neurologically intact individuals – exhibit imperfect somatosensory control of the arm and hand in the absence of ongoing visual feedback (Fuentes and Bastian, 2010; Paillard and Brouchon, 1968; Smeets et al., 2006; Wann and Ibrahim, 1992) The most conspicuous and ubiquitous manifestation of imperfect somatosensation is “proprioceptive drift” (Wann and Ibrahim, 1992; c.f. Smeets et al., 2006), wherein bias in the perceived position of the unseen hand develops within a period of 12 to 15 seconds (a progressive degradation in the accuracy of proprioceptive sensation; c.f., Paillard and Brouchon, 1968). A second manifestation is a lack of precision in the proprioceptive estimation of limb state, whereby repeated estimates of hand position or joint configuration exhibit marked variability (c.f., Fuentes and Bastian, 2010).

We devised a planar reaching task wherein neurologically intact people grasped the handle of a 2-joint robotic manipulandum while making goal-directed movements between 25 spatial targets arranged in a 5 × 5 grid. To differentiate between the effects of short-term training with supplemental limb state feedback on subsequent performance of reaching movements made both with and without that feedback, we designed the inter-target distance of this grid to be less than the magnitude of spatial uncertainty in the proprioceptive assessment of hand position as reported by Fuentes and Bastian (Fuentes and Bastian, 2010). At the same time, we implemented a vibrotactile display of the arm’s planar work space wherein a hand displacement corresponding to the grid’s minimum inter-target distance also corresponded to a change in vibrotactile stimulation that was greater than three times the just-noticeable differences in the magnitude of that stimulus, as reported by Shah and colleagues (Shah et al., 2016). These design choices enabled us to test two main hypotheses: (1) that neurologically-intact humans can learn to use vibrotactile limb state feedback to enhance the on-going control of a moving limb beyond the limits of intrinsic proprioception; and (2) that this training has aftereffects on subsequent movements performed without concurrent supplemental vibrotactile feedback (i.e., that training induces improved proprioceptive control; c.f., Cuppone et al., 2016). Preliminary aspects of this study have been presented in abstract form (Risi et al. 2016, 2017).

## Methods

### Subjects

Fifteen right-handed, neurologically-intact human adults [6 female; 23.8 ± 4.1 years (mean ± SD, here and elsewhere)] were recruited from the Marquette University community. All provided written informed consent to participate in the experimental procedures of this study, which were approved by Marquette University’s Office of Research Compliance. None of the subjects had known neurological disorders, all had normal or corrected-to-normal vision, and all were naive to the purposes of the study.

### Experimental setup

Each subject was seated in a high-backed, adjustable-height chair that was placed in front of a horizontal planar robotic manipulandum (Fig 1A; c.f., Scheidt et al. 2010). Subjects grasped the robot’s handle with their right hand. The left arm rested comfortably in a cloth sling with the forearm and hand pointed forward. A horizontal opaque shield covered the workspace to block the subject’s view of their moving arm and the robotic apparatus. View of the stationary arm was not precluded. Seat height was adjusted such that the shoulder was aligned with the opaque shield, thus allowing the subject to comfortably move the robot’s handle throughout the workspace. To display visual feedback, a 42” computer screen was oriented vertically above the shield within direct view (0.7 m in front of the subject). This screen provided real time visual cues of hand and target positions when appropriate (the visual feedback schedule is described below).

**Figure 1:**
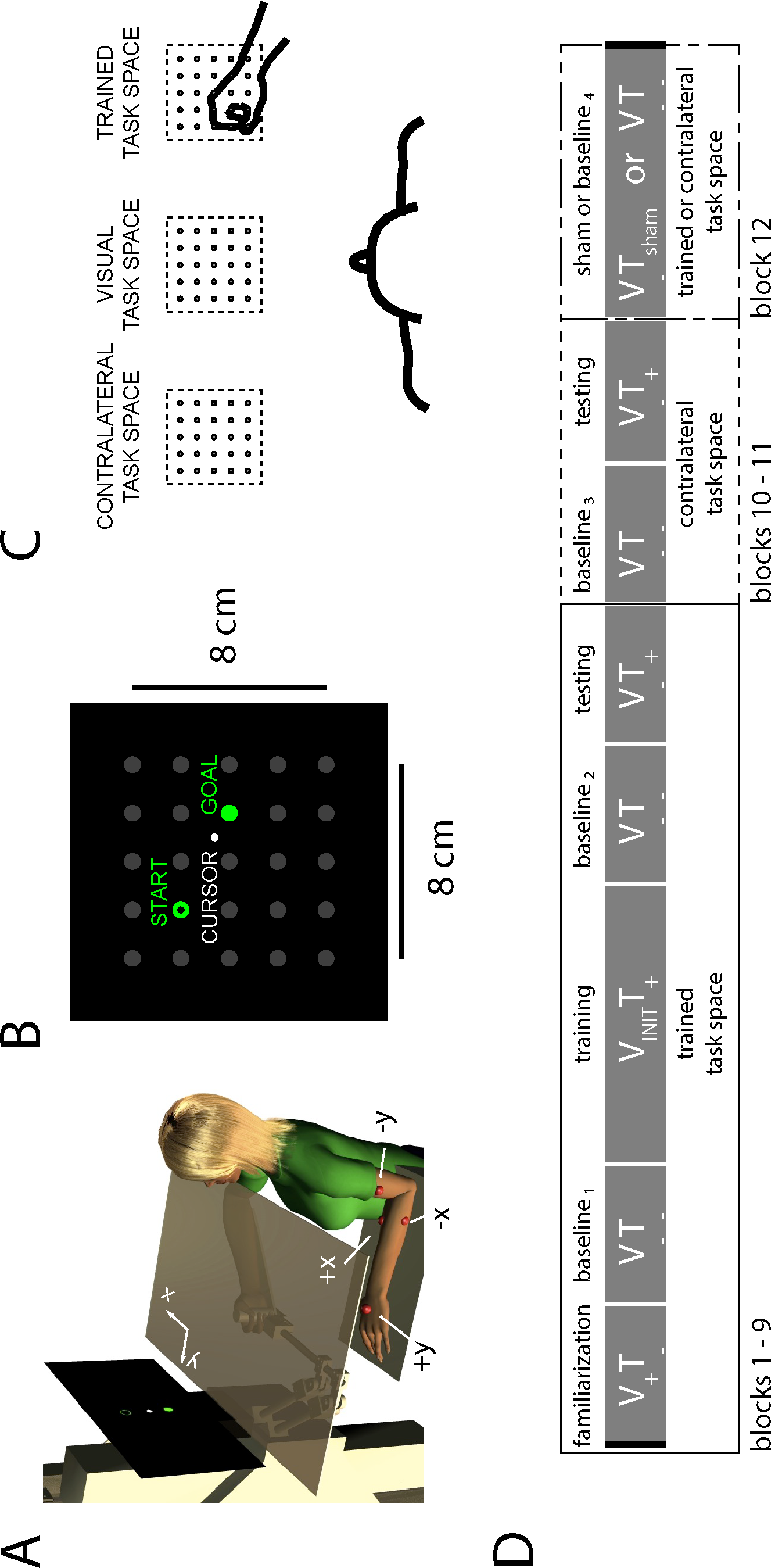
Experimental setup and protocol. A) Subject holding the end effector of a planar manipulandum, with visual occlusion shield. Red spheres indicate default configuration of the vibrotactile display. B) Displayed grid and visual cues (cursor and targets). C) The visual display was aligned with the subject’s midline. The subject was tested in both the right task space (i.e. trained task space) and the left task space (i.e. untrained contralateral task space). D) Sequence of experimental blocks. The familiarization, baseline, testing, and sham blocks each consisted of 25 reaches. The training block consisted of 5 sets of 25 reaches. Visual feedback (V) and vibrotactile feedback (T) of hand position were either continuous (+), absent (−), or only displayed briefly prior to the onset of the reach (init). Subjects performed two experimental sessions on two separate days.

Supplemental vibrotactile feedback of the moving hand’s position was provided using a two-channel (4 tactors) vibrotactile display attached to the non-moving arm. The tactors (Pico Vibe 10 mm vibration motors; Model # 310-117; Precision Microdrives, Inc,) have an operational frequency range of 50 to 250 Hz and peak vibrational amplitude of 0.97 N, which corresponds to an expected maximal forearm-plus-hand acceleration ranging between 0.53 m/s^2^ and 0.77 m/s^2^, depending on subject anthropometrics. Tactors were taped to the skin over four different dermatomes (c.f., Lee et al. 2008) with default locations indicated by red spheres in Fig 1A. Two tactors encoded hand position along the x-axis of the workspace whereas the other two tactors encoded hand position along the y-axis. The +y tactor was placed on the skin above the ulnar head (dermatome C8); +x tactor was placed above the flexor carpi radialis muscle (dermatome T1); the −y tactor was placed above the brachialis muscle (dermatome C5); the −x tactor was placed above the extensor carpi ulnaris muscle (dermatome C7). Elastic fabric bands were used to secure the tactors. The vibrotactile encoding scheme was designed such that motions of the robot handle to the right induced the +x tactor to vibrate whereas motions of the robot handle away from the subject (i.e., toward the monitor) induced the +y tactor to vibrate. A detailed description of the vibrotactile encoding scheme is provided below. For additional details on tactor placement and their computer-controlled activation see Krueger et al. (2017).

### Task space and vibrotactile display

Subjects performed reaches between 25 targets (0.6 cm radius dots) arranged in a 5×5 grid that had an edge length of 8 cm (Fig 1B). We selected the minimum inter-target distance (2.0 cm) to be smaller than the range of uncertainty of proprioceptive perception of limb position (2.5 ± 0.2 cm) as derived using joint angular uncertainty values reported by (Fuentes and Bastian, 2010). We constructed a mapping between the task space and its representation within the vibrotactile display such that very small displacements of the hand were clearly discriminable within the vibrotactile display (Fig 2). Under this mapping, the just noticeable difference (JND) in magnitude of sequential vibrotactile stimulations as reported by (Shah et al. 2016) (i.e., a change of 16.3 ± 11.8 Hz) corresponded to 0.3 ± 0.2 cm of hand displacement. Therefore, the minimum inter-target distance was more than 6 times greater than the vibroctatile JND and considerably smaller than the range of uncertainty of proprioceptive perception of limb position.

**Figure 2:**
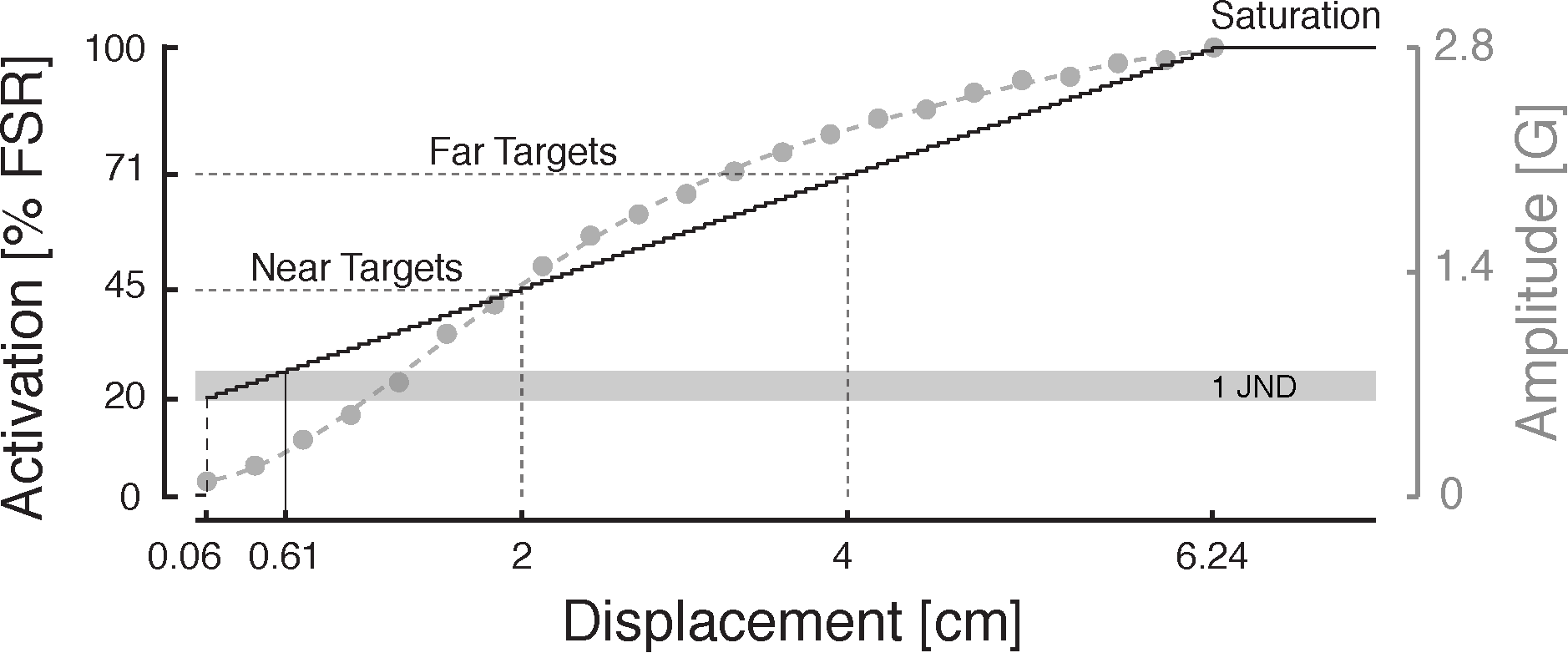
Vibrotactile encoding scheme: *position* feedback. Left axis: activation of one tactor (Percentage of Full Scale Range) plotted against handle displacement along the direction encoded by the tactor. The height of the gray box indicates the average discrimination threshold for dermatome C7 (from Shah et al. 2016). Right axis: tactor vibration amplitude is a monotonic function of hand displacement along the encoding direction.

Specifically, we employed a piecewise-linear activation map (Fig 2) wherein tactor activation intensity was 71% full scale range (FSR) (i.e., 3.5 V) when the hand was on a far target (4 cm from the center of the task space), 45% when the hand was on a near target (2 cm from the center), and 26% when the hand was on the outer edge of the central target (i.e., 0.6 cm from its center). The feedback was turned off in all tactors (i.e. the activation was 0.0 V) only when the hand was centered on the central target (i.e., within 0.06 cm from the grid center). This range of activations corresponded to tactor vibration amplitudes ranging from 0 G to 2.0 G. The discontinuity at 0.06 cm served to counteract static friction within the tactor motor. Within the task space, the intensity of vibration increased in the vibrotactile display as a vector representation of the hand’s deviation from the center target position. The frequency of vibration (in units of Hz) was roughly 100 times the amplitude of vibration (in units of G).

### Experimental Protocol

Subjects performed two experimental sessions on separate days within two weeks (range: 1 to 10 days). Prior to testing on each day, subjects were introduced to the vibrotactile display and were invited to freely explore the robot’s workspace by displacing the robot’s handle along the two cardinal axes. During this period, subjects were frequently and repeatedly asked to report which tactors were activated at any given time. If subjects made errors in detecting vibration on any tactor, we adjusted that tactor’s location by 2 or 3 cm so that each subject could reliably detect and report vibration. Subjects were then encouraged to explore the vibrotactile display by making self-guided reaching movements. This introduction and exploration procedure took between 2 to 5 minutes to complete. Two to five minutes of exploration were sufficient to provide a basic understanding of how a portion of the arm’s larger workspace was encoded within the vibrotactile feedback.

During the main part of both experimental sessions, subjects performed 12 blocks of 25 reaching movements, one reach per trial. The target grid was always displayed on-screen in low-contrast gray, with the current target presented in vivid green (Fig 1B). Subjects were instructed to “capture the target as accurately as possible”. Upon completing a reach, subjects announced that they had arrived at the target and the experimenter registered the event by pressing a button. Subjects had 10 seconds to complete each reach. At the end of the trial, to emphasize the beginning of a new trial, the previous target became an empty green dot and, after few seconds (an inter-trial interval of 2.3 ± 0.7 sec), a different dot turned green providing the cue to move. The target sequences were pseudo-randomized across each block of 25 trials, with the distance between consecutive targets in the range 4.0 to 6.32 cm.

The sequence of trial blocks in the two experimental sessions were identical except for the last block. The first nine blocks were performed with the center of the physical target grid shifted 20 cm to the right of the subject’s midline (i.e., trained task space, Fig 1C). As all subjects used their right hand to move the robotic handle, subjects practiced primarily on the ipsilateral side of the arm’s reachable workspace. Subjects then performed two test blocks with the center of the physical target grid shifted 20 cm to the left of the subject’s midline (i.e., on the contralateral side of the arm’s workspace, i.e., in the contralateral task space; Fig 1C). The last block was performed in either the trained task space (Day 1) or the contralateral task space (Day 2). By contrast, the center of the visual grid was aligned at all times with the subject’s midline (i.e., *visual task space*, Fig 1C).

The 12 blocks were performed under various combinations of visual (V) and vibrotactile (T) feedback of hand position (Fig 1D). Visual and vibrotactile feedback were either continuous (+), absent (−), or only displayed briefly prior to the onset of the reach (INIT). During the first trial block (Fig 1D, *familiarization*, trained task space), subjects were familiarized on the task by performing 25 reaches with visual feedback but without vibrotactile feedback (V_+_T_−_); a small white visual cursor (0.5 cm radius) continuously tracked hand position during this block. Next, in order to assess baseline accuracy before training with supplemental vibrotactile feedback, subjects performed a block of reaches without any performance feedback (V_−_T_−_; Fig 1D: *baseline 1*, trained task space). Five blocks of training with vibrotactile feedback followed the baseline assessment; here, the visual feedback of the cursor was limited only to the moment just prior the reach onset (V_INIT_ T_+_; Fig 1D: *training*, trained task space). Throughout these movements, vibrotactile feedback of the state of the moving right arm was provided to the stationary left arm, whereas visual feedback of the cursor was provided only briefly when the cursor was inside the starting target, disappearing when the new target appeared. No cursor or other visual cues were provided during training trials in order to minimize visual distraction from the vibrotactile feedback. During training, subjects were also provided haptic knowledge of results in the form of robot-guided corrections to target capture errors. That is, upon completing each training reach (i.e., when the experimenter pressed the button indicating target capture or after the 10 second timeout), the desired target turned white, while the robot gently guided the subject’s hand to the correct target location (unless the subject had correctly hit the target). Upon completion of the robotic correction, a new desired target turned green. On average, the five blocks of training lasted 30 minutes in total.

We then examined aftereffects of training by assessing post-training baseline performance without either visual or supplemental vibrotactile feedback (V_−_T_−_; Fig 1D: *baseline 2*, trained task space). This block quantified both the extent to which performance improvements observed during training were due specifically to the presence of concurrent vibrotactile feedback as well as the magnitude of performance gains induced by an improved ability to use intrinsic proprioception to guide reaching movements (i.e., perceptual learning). We then tested the extent to which subjects could recall what they had learned during training by having them complete one test block of reaches with only vibrotactile feedback and without visual feedback or haptic corrections (V_−_T_+_; Fig 1D: *testing*, trained task space).

Next, we assessed the capability of subjects to apply what they had learned in the trained task space to reaches performed in the contralateral task space. Here, a baseline block without visual or vibrotactile feedback was performed in a physical task space centered 20 cm to the left of the subject’s midline (V_−_T_−_; Fig 1D: *baseline 3*, contralateral task space). This block was followed by a “contralateral task space test block” of reaches with only vibrotactile feedback and no visual feedback (V_−_T_+_; Fig 1D: *testing*, contralateral task space).

Finally, the last 25 reaches were performed with different feedback conditions in the two experimental sessions. On Day 1, subjects reached in the ipsilateral (right) task space guided by sham vibrotactile feedback (V_−_T_SHAM_; Fig 1D: *sham,* trained task space). In this feedback condition, the vibration recorded during a training block was played-back without reference to the subject’s actual performance. Providing non-informative vibrotactile feedback allowed us to assess whether potential performance enhancements were due specifically to the information contained within the supplemental vibrotactile feedback and not by the mere presence of vibration (see also Krueger et al. 2017). Subjects who noticed that the vibration lost its information content were instructed to nevertheless attempt to use the “feedback” as best they could; we informed subjects only after the end of the experimental session that this last block involved non-informative vibration. During the second experimental session, the last block was meant to assess potential order effects of test blocks in the contralateral task space, i.e. the extent to which test block performance improvements in the contralateral task space might simply have been due to additional practice on the task during the post-training baseline. Therefore, instead of sham feedback, subjects performed an additional 25 baseline reaches in the contralateral task space without either visual or vibrotactile feedback (V_−_T_−_; Fig 1D: baseline 4, contralateral task space). At the end of both sessions, subjects were asked to report how they used the information encoded in the vibratory feedback to guide their reaching movements.

### Data Analysis

Analysis of kinematic performance focused principally on hand position at two time points during each reach, i.e. when the target first appeared (*initial hand position*) and when the subject indicated that they had acquired the intended target (*final hand position*). Our primary measure of accuracy was the *target capture error*, which is defined as the absolute distance between the final hand position and the target location.

We derived eight additional task-related variables. We defined the *actual movement vector* as a vector pointing from the initial to the final hand position [Fig 3A, (1)]. We defined the *ideal movement vector* as the vector pointing from the actual initial hand position to the intended target location [Fig 3A, (2)]. We defined *directional error* β (Fig 3A) as the magnitude of the angle between the actual movement direction (solid black line) and the ideal movement direction (dashed black line) (see Gordon et al. 1995, Ghez et al. 1995). β values are small if the subject correctly estimates his/her initial hand position and moves the hand in the proper direction (cf., Fig 3A). We defined *extent error* as the difference between the actual movement magnitude and the ideal movement magnitude; extent error is positive if the actual movement extent exceeds the ideal extent (Gordon et al. 1995, Ghez et al. 1995).

**Figure 3:**
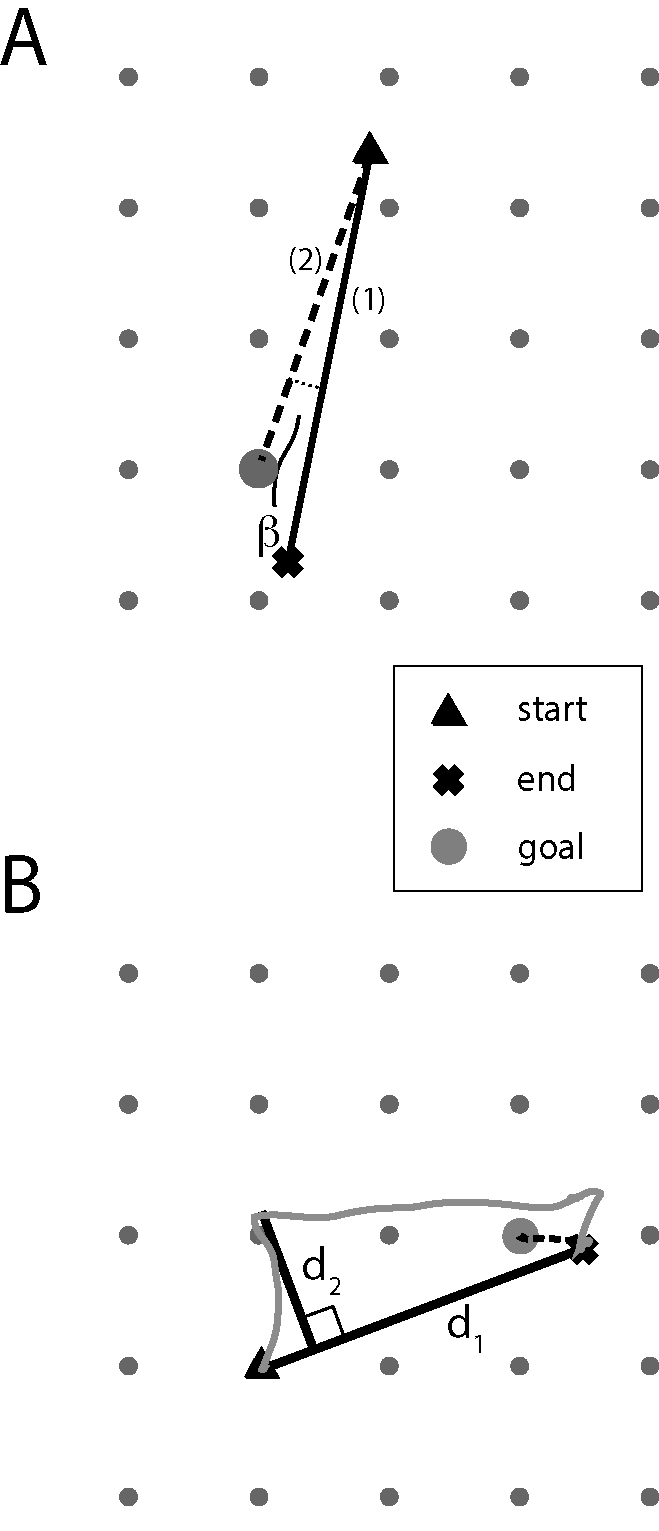
Single trial performance in two distinct trials. Triangles: initial hand position; Large gray circle: current intended target location; Cross: final hand position. A) Depiction of direction and extent error; the black dashed line shows the ideal movement direction and extent; the black solid line shows the actual movement direction and extent. Movement extent error is defined as the length of line (2) subtracted from the length of line (1). *β* is defined as the angle between the ideal and actual movement vectors. The angle is positive if the rotation from the actual vector (solid line) to the ideal vector (dashed line) was counter clockwise. B) Depiction of aspect ratio calculation. Gray path: a subject’s hand trajectory. Black solid lines: d1 is the segment between the initial position and the final position; d2: the perpendicular distance between line segment d1 and the point of greatest hand path deviation from d1.

We computed the *spatial contraction/expansion* of reach endpoints along both coordinates (i.e., *cont*/*exp*_*xy*_) to assess the fidelity of the subject’s internal representation of extrapersonal space in both task spaces (c.f., Dukelow et al. 2010). For the outer targets, spatial contraction/expansion was computed as the area of the quadrilateral spanned by the final hand positions of reaches made to the outer targets of the grid. This area was then normalized to (i.e., divided by) the actual area of the outer grid square. Spatial contraction/expansion was similarly computed for movements made to the inner targets of the grid.

We also evaluated the extent to which subjects’ trajectories became straighter as they practiced using the vibrotactile feedback. To this end, we defined *aspect ratio* as the ratio |*d2*|*/*|*d1*|, where |d*1*| was defined as the distance between the initial hand position and its final position, and |*d2*| was defined as the perpendicular distance between line segment d1 and the point of greatest hand path deviation from d1 (Fig 3B). The aspect ratio is a larger number (closer to one) when the subjects used an inefficient path and is a small number when the subjects used nearly a straight-line path. We also introduce a new scalar performance measure-the *decomposition index, DI* (Eqn 1) – a unitless performance measure that together with aspect ratio, can distinguish between different strategies for using vibrotactile feedback to solve the target capture task in the absence of visual feedback. Specifically, the DI quantifies the extent to which sampled-data hand paths in any given trial move exclusively parallel to the cardinal {x, y} axes of the vibrotactile display:

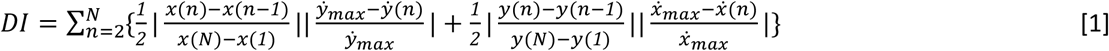

where N corresponds to the maximum number of data samples within a given trajectory, n corresponds to the sample number in that trajectory, whereas *ẋ*_*max*_, and *ẏ*_*max*_, correspond to the peak hand speeds along each of the cardinal axes. If the hand first moves parallel to the x axis (i.e., y(n) − y(n − 1) = 0) and then parallel the y-axis (x(n) − x(n − 1) = 0), the initial motion along the x axis will yield zero values of *ẏ*(*n*), and thus the equation will “integrate” the x-axis displacements with unitary weight in the first half of the movement. In the second half of this hypothetical movement, displacements along the y-axis are integrated with unitary weight because the x-axis velocity is zero during this portion of the trajectory. Thus, the overall DI value would be high. By contrast, if a movement were to be directed along a diagonal or any off-axis movement, the largest sample displacements along the x-axis would occur at the same time as the peak speed along the y-axis leading to small contributions to DI; a similar argument holds for the y-axis displacements. In this second hypothetical case, the DI would be small. As presented in Appendix I, off-axis, straight-line trajectories with bell-shaped velocity profiles yield DI values equal to 0.24.

Finally, we computed *target capture time* as the difference between the moment when the target appeared and the moment when the subject indicated she/he had acquired the intended target.

### Statistical hypothesis testing

This study tested two main hypotheses. The first proposes that neurologically-intact subjects can learn to use supplemental vibrotactile feedback of hand position to enhance the ongoing control of reaching movements beyond limits imposed by intrinsic proprioception. The second proposes that this training has aftereffects on subsequent movements performed without concurrent supplemental limb state feedback (i.e., that training induces improved proprioceptive control; c.f., Cuppone et al., 2016). To test these hypotheses, we compared target capture error across feedback conditions and days.

More specifically, we designed the sequence of training blocks to facilitate planned paired t-test comparisons between specific training blocks (feedback conditions) either within or across days. To test for the presence of sensorimotor learning (the first hypothesis), we compared target capture errors in the initial training block to those in the final training block in each day. We also compared reach performance driven by intrinsic proprioceptive feedback alone against performances driven additionally by vibrotactile limb state feedback by comparing target capture errors in the two post-training test blocks in the trained task space (i.e., comparing blind reach performance with and without vibrotactile feedback). These analyses were performed separately for the training blocks of day 1 and day 2. To test for aftereffects of vibrotactile feedback training on subsequent movements performed without either visual or vibrotactile feedback (i.e., the second hypothesis pertaining to perceptual learning), we compared target capture errors between no-vibration baseline blocks performed immediately before and after the five training blocks on both days.

We performed ancillary paired t-test comparisons to examine the specificity of action for sensory augmentation using vibrotactile feedback. We used paired t-test to compare target capture errors across test phase blocks with and without vibration in the contralateral (untrained) task space to test if the ability to use vibrotactile state feedback extends beyond the trained workspace. We compared the V_−_T_+_ condition verses V_−_T_SHAM_ condition on Day 1 to verify that any performance improvement observed when using supplemental feedback was due specifically to the information encoded within the vibrotactile display, rather than to the mere presence of vibration. We also tested for retention of learning across days by comparing target capture errors within the initial training sets on days 1 and 2. We also compared Day 2 baseline block performance within the contralateral task space immediately before and after the test block with vibration (i.e., block 9) to verify that any potential benefits of vibration in the contralateral task space were indeed due to the vibrotactile feedback, rather than being due to a possible order effect.

Additional supplemental statistical tests were performed on the secondary performance measures (spatial contraction/expansion, extent and directional errors, decomposition index, target capture time) to address specific questions related to the strategies used by subjects to solve the target capture task using vibrotactile feedback. All statistical tests were carried out in the SPSS computing environment (SPSS version 24.0; IBM Corp.). We report results of the ancillary analyses after Bonferroni-correction such that effects were considered statistically significant at the α = 0.05 level.

## Results

This study sought to characterize how human subjects learn to use supplemental vibrotactile feedback of performance to enhance reach accuracy beyond limits imposed by the inherent inaccuracies of intrinsic proprioception, as reported in the literature (c.f., Fuentes and Bastian, 2010). We asked subjects to perform a horizontal planar reaching task in two task spaces. We constructed a mapping between the position of the moving hand and its representation within the vibrotactile display such that small, goal-directed hand displacements were more easily discriminable using supplemental vibrotactile feedback than with intrinsic proprioceptive feedback in the moving arm. Subjects then made goal-directed (cued) reaches to specific locations within the task spaces.

### Target Capture Error

Figure 4 depicts all final end-of-reach hand positions (gray dots) achieved on Day 1 by a selected subject; this performance was typical of the population (see group data reported below and in Fig. 5). During the familiarization block when visual cues were provided (Fig 4: *familiarization*), the subject was able to accurately reach all targets (i.e., the reach endpoints sampled the target task space uniformly; shown with the gray shaded region). Target capture performance degraded dramatically when the visual cues were subsequently removed (Fig 4: *baseline 1*, trained task space): the subject exhibited both a systematic drift towards the body midline as well as a spatial expansion of the performance space.

**Figure 4:**
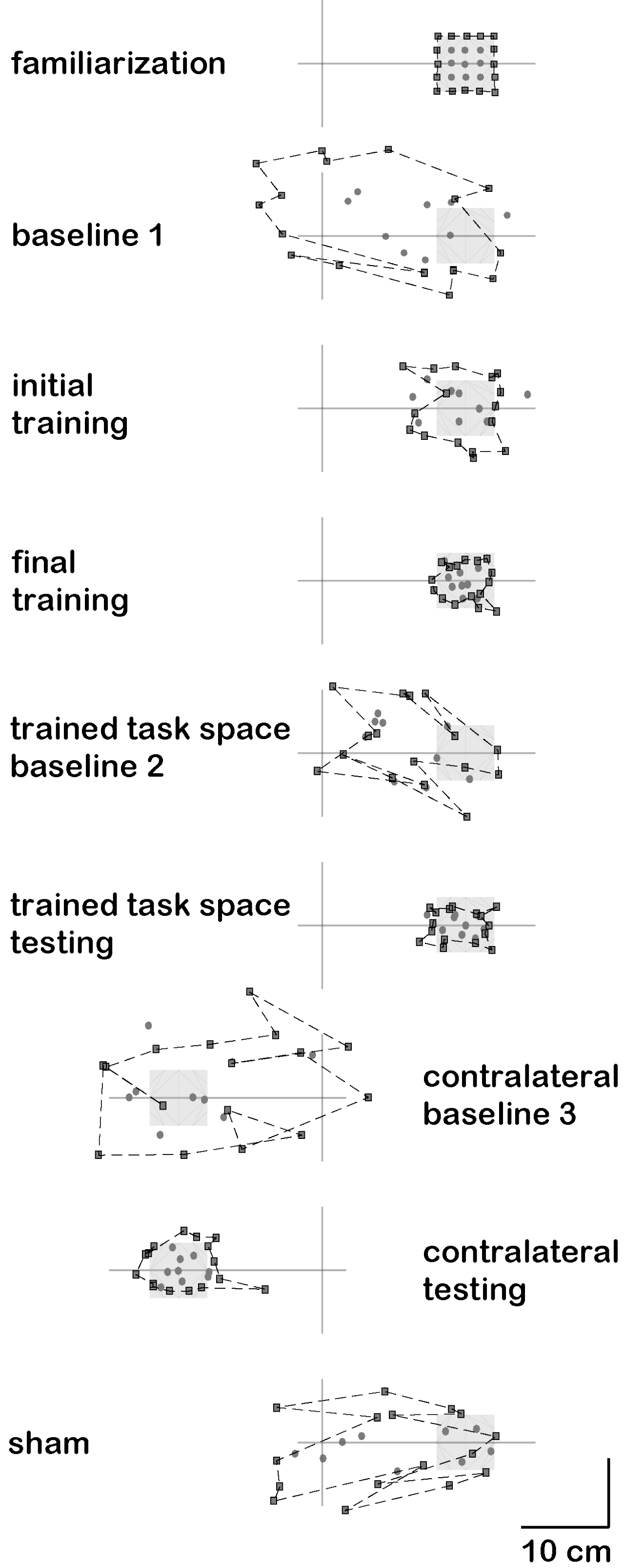
Performance of a selected subject in the reaching task on Day 1. For each of the blocks, hand positions after reaching are plotted for inner targets (circles) and outer targets (squares). The spatial location of the target grid and its location relative to the center of the visual workspace (axis lines) is represented by the shaded square. A dashed line connects the outer targets. The progression of experimental blocks ranges from top to bottom in the figure.

**Figure 5:**
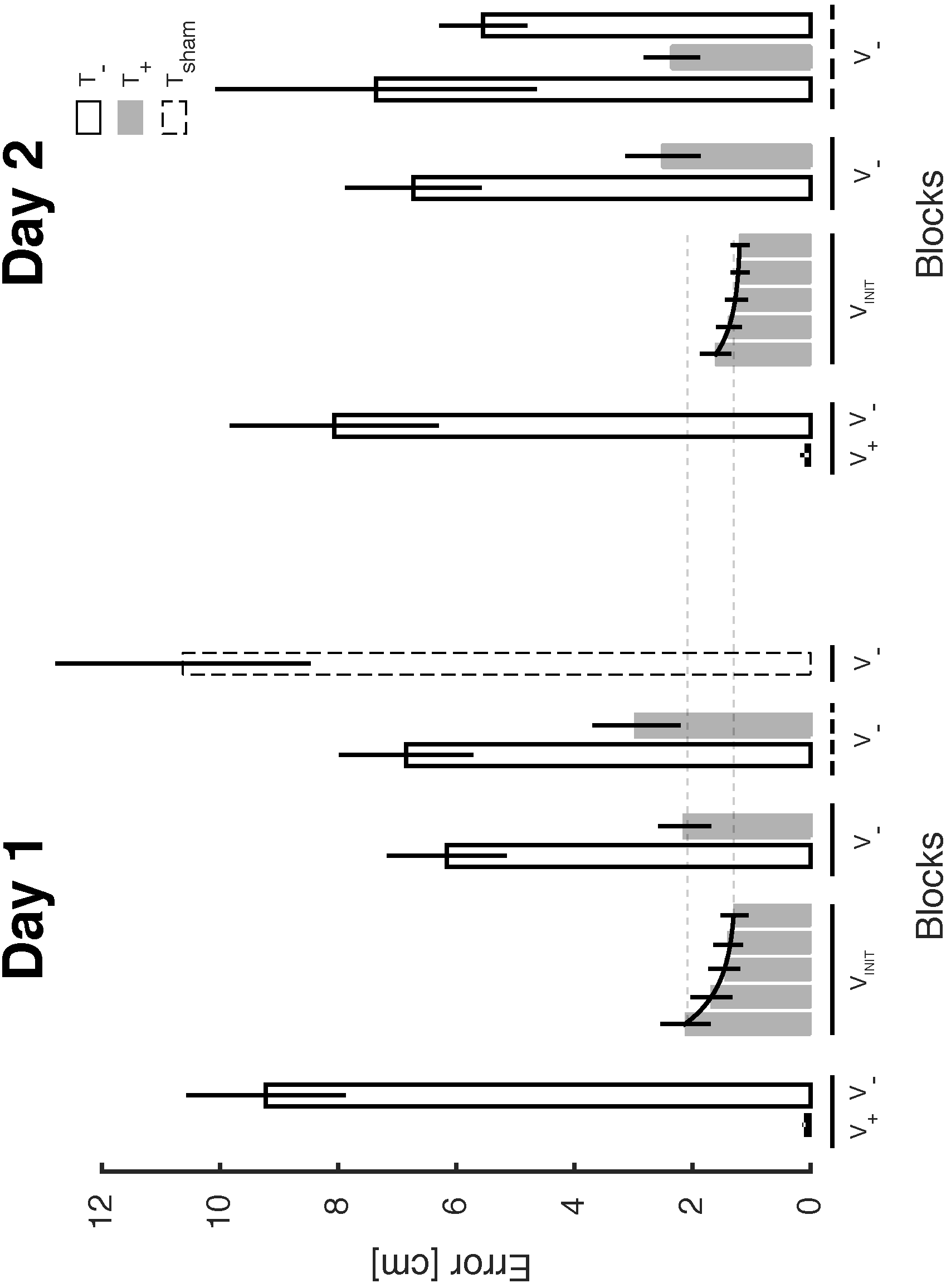
Group results: Root Mean Square (RMS) target capture error vs. experimental block for both days. Black bars: the initial block with visual feedback (V_+_) at the start of each experimental session; V− indicates blocks without visual feedback and V_INIT_ indicates visual feedback of the cursor only at the beginning of a trial while the subject was near the starting target. Gray bars: blocks with vibrotactile feedback (T_+_); White bars: blocks without vibrotactile feedback (T_−_); Dashed bar: sham feedback (T_sham_). Horizontal solid bars under the x-axis indicate blocks in the training workspace while the horizontal dashed lines indicate blocks in the generalization workspace. Note that we did not compare target capture errors across the T_+_ blocks in the V_INIT_ and V_−_ conditions because differences in the visual feedback conditions at the onset of training trials can differentially affect the accumulation of proprioceptive drift (c.f., Wann and Ibrahim, 1992). Error bars: 95% confidence interval (CI) about the mean.

Next, the subject performed five blocks of training with vibrotactile limb state feedback (approximately 30 minutes); although subjects were not provided concurrent visual feedback of cursor motion, they were provided knowledge of results in the form of robot-guided corrections at the end of each reach. These terminal corrections were intended to prevent the accumulation of proprioceptive drift. Across the five training blocks on Day 1, target capture errors decreased (accuracy improved) from the initial to the final block of training (Fig 4: *initial training*, *final training* respectively).

After training, the subject exhibited a reduced expansion of the task space during post-training baseline assessment (Fig 4: trained task space *baseline 2*), thus demonstrating a beneficial aftereffect of training, even without concurrent vibrotactile feedback. By contrast, proprioceptive drift towards the midline did reappear during the post-training baseline 2 block. When vibrotactile feedback was reinstated, the drift immediately resolved (Fig 4: trained task space *testing*). A similar pattern of behavior arose when the subject was tested in the contralateral task space: we observed drift toward the body midline and spatial expansion in the absence of vibrotactile feedback (Fig 4: contralateral baseline 3). With concurrent vibrotactile feedback, there was an increased target capture accuracy and reduced proprioceptive drift (Fig 4: contralateral testing). Target capture performance subsequently degraded when non-informative feedback was provided in the trained task space (Fig 4, compare *sham* to *trained task space testing*). These results demonstrate that the performance enhancements seen with supplemental limb state feedback were specifically due to the information encoded within the vibrotactile display, and not simply due to the mere presence of vibration applied to the non-moving arm.

The individual-subject results were characteristic of the study cohort as a whole. Across the study population, subjects reached the targets with accuracy and precision in the presence of concurrent visual feedback whereas performance degraded markedly when visual cues were subsequently precluded and no supplemental feedback was provided (Fig 5: compare the two left-most bars in each panel). Remarkably, reach errors dropped precipitously and immediately upon providing vibrotactile feedback (compare the second to the third left-most bars in Fig 5). To characterize the extent to which subjects might improve their ability to use vibrotactile limb state feedback, we examined differences in accuracy across the five training blocks (i.e., the five shaded bars above the horizontal bar marking the V_INIT_ trials). On Day 1, errors decreased most rapidly within the first two to three blocks of training. Target capture errors appeared to reach a performance plateau by the third block of training, with little further improvement in performance in subsequent training blocks. Similar patterns of performance changes were observed across trial blocks in both experimental sessions.

We first challenged the hypothesis that neurologically-intact subjects could learn to use vibrotactile limb state feedback to enhance the on-going control of reaching movements beyond limits imposed by intrinsic proprioception. One-sided paired t-test found within-day reduction of target capture errors during training to be significant on both days (T_14_ > 3.99, p < 0.0005 in both cases). Performances immediately and similarly degraded when vibration was removed after training (i.e., when subjects had to rely solely on intrinsic proprioceptors to perform the task in the Fig 4 *baseline 2* block and in Fig 5, the first white bar to the right of the training bars). During post-training *testing* blocks, the performance enhancing effects of supplemental vibrotactile feedback were immediately restored (c.f., Fig 5: gray bars above solid horizontal V_−_ lines) such that target capture errors dramatically and significantly decreased relative to the post-training baseline block (one-sided paired t-test: T_14_ > 6.98; p < 0.0005 on both days). These findings confirm that subjects could learn to use vibrotactile limb state feedback to improve performance in the trained task space beyond levels typically observed when using intrinsic proprioception alone.

Across the five blocks of training on each day, the learning trend was reasonably well characterized by a falling exponential decay of performance error, with learning rate time constants averaging 1.4 ± 1.1 blocks on Day 1 and 1.3 ± 0.9 blocks on Day 2. We found evidence that RMS target capture error decreased between the initial training block of Day 1 to the initial training block of Day 2 (one-sided paired t-test: T_14_ = 2.25, p < 0.05), by which we infer that subjects retained knowledge gained through training from Day 1 to Day 2. This increased accuracy was due to the specific information encoded within the vibrotactile display – and not merely the vibratory stimulation itself-because performance degraded once again when non-informative sham feedback was provided at the end of Day 1 (Fig 5 left: dashed bar; one-sided paired t-test between post-training recall and sham: T_14_ = 6.26, p < 0.0005).

We next tested that the proposed training has aftereffects on subsequent movements performed without such feedback. Consistent with this premise, when vibrotactile feedback was removed after training on Day 1, target capture errors in the baseline-2 V_−_T_−_ block (Fig 5: first white bar to the right of the training bars) were substantially lower than those in the baseline-1 V_−_T_−_ block (Fig 5: second white bar from the left) one-sided paired t-test: T_14_ = 4.1778, p < 0.0005). This beneficial aftereffect of training was evidently limited to the initial bout of training, as no further improvements were observed after a second bout of training (i.e., when comparing baseline-2 performance on Day 2 to baseline-2 performance on Day 1: T_14_ = 1.1140, p = 0.284).

Consistent with the hypothesis that the utility of vibrotactile limb state feedback generalizes to movements performed in untrained regions of the reachable workspace, we observed immediate performance enhancements due to the vibrotactile feedback also in the contralateral workspace. Specifically, we observed a marked improvement in reach accuracy between the *baseline-3* and *testing* blocks in the contralateral untrained workspace (Fig 5, white vs gray bars above dashed, bottom line) (one-sided paired t-test: T_14_ = 7.15, p < 0.0005). The observed benefits of vibrotactile feedback were not simply due to an order effect, as confirmed by the baseline-3 block (V_−_) at the end of the Day 2 (Fig 5, second white bar above dashed, bottom line in right panel), wherein RMS errors increased relative to the prior testing block (one-sided paired t-test: T_14_ = 7.64, p < 0.0005). We observed no noticeable differences in target capture error within the same testing conditions across the two task spaces within each day. These results support the hypothesis that the benefits of supplemental vibrotactile feedback training generalize across the arm’s reachable workspace.

### Direction and Extent Errors

We analyzed directional and extent errors (see Fig 3 for a description) to gain insight into the effect of concurrent vibrotactile feedback on proprioceptive drift. Because even a glimpse of hand position feedback can arrest or eliminate drift (Wann and Ibrahim, 1992), we present in Fig 6 only those blocks performed entirely without visual cues (pre-training: *baseline 1*; post-training: *baseline 2* and *testing* in trained task space, and *baseline 3* and *testing* in the contralateral task space). Whereas unsigned directional error varied markedly across experimental blocks, we observed no evident effect of days, nor any apparent interaction between experimental blocks and days. On both days, directional error decreased in the trained workspace from *baseline 1* and *baseline 2* (no vibrotactile limb state feedback) to *testing* (with vibrotactile feedback) (Fig 6A white bars vs. gray bars with solid lines below). A similar trend was reported in the contralateral workspace (Fig 6A white vs gray bars with subjacent dashed lines). Movement extent errors similarly varied across experimental blocks (Fig 6B), with the most striking and consistent differences observed between *baseline* (no vibrotactile feedback) and *testing* (with vibrotactile feedback) in both the trained and generalization task spaces (Fig 6B; white vs. gray bars to the right of the vertical dotted line in each day). Thus, we infer that subjects used limb state information encoded within the supplemental vibrotactile feedback signals to correct their initial drift and to improve movement accuracy.

**Figure 6:**
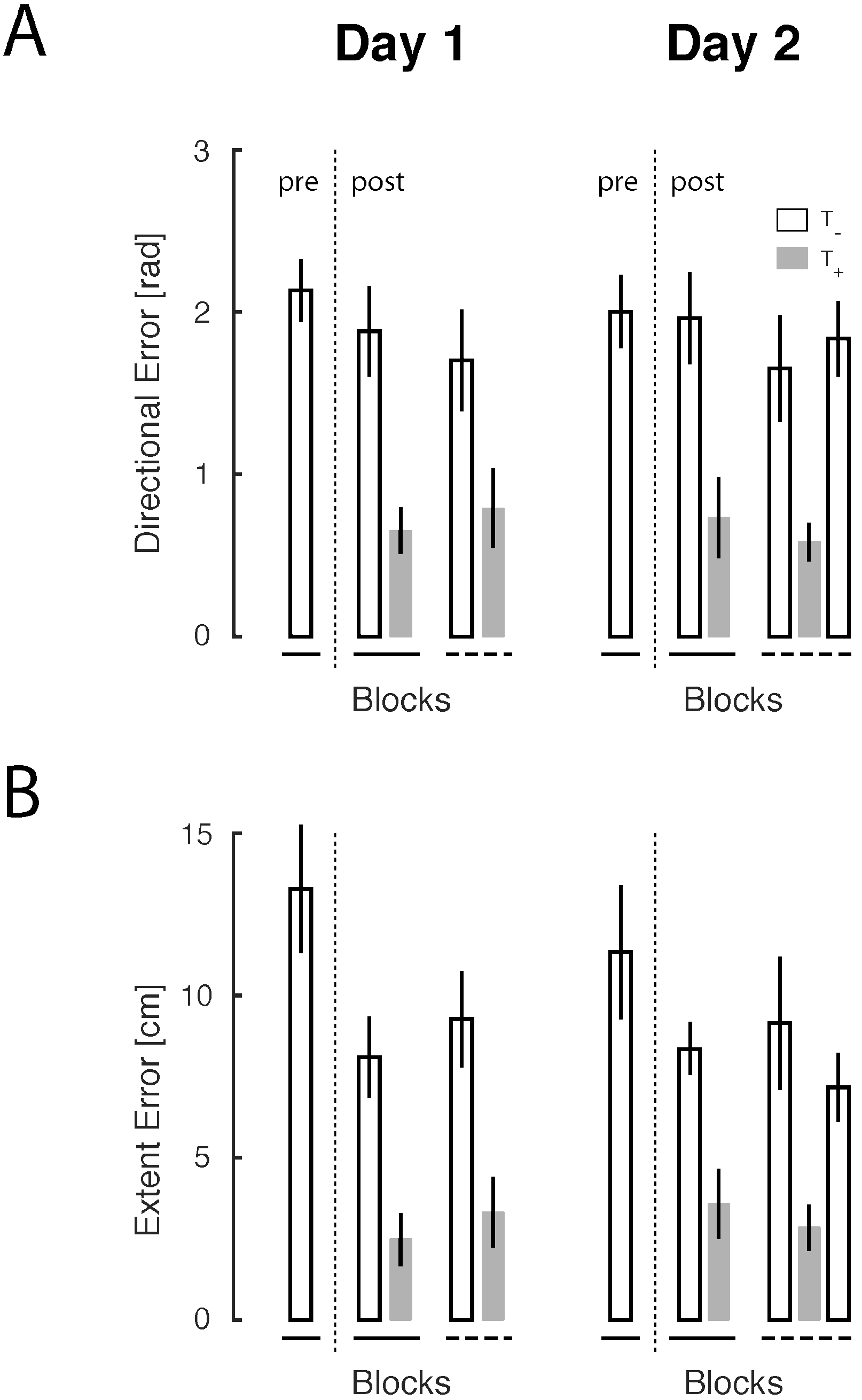
Group results for direction and extent errors across blocks for both days. **A)** Direction error vs. blocks in experimental phases wherein visual cues were precluded (V_−_). Pre-training: *baseline 1*; post-training: *baseline 2* and *testing* in the trained workspace (white and gray bars above solid horizontal line), *baseline 3* and *testing* in the generalization workspace (white and gray bars above dashed horizontal line). The vertical dotted line indicates where training occurred in the sequence of trial blocks. **B)** Extent error vs. blocks in experimental phases wherein visual cues were precluded. The presentation of trial block performance is as described for panel (A). Error bars: 95% CI; Gray bars: blocks with vibrotactile feedback (T_+_); White bars: blocks without vibrotactile feedback (T_−_).

### Spatial Contraction/Expansion

The analysis of spatial contraction/expansion (*cont/exp_xy_*) revealed an expansion of target space when visual cues were not provided (Fig 7). However, the *cont/exp_xy_* index was closer to the ideal value of 1.0 when the vibrational feedback was provided in *testing* blocks relative to *baseline* blocks, both in the trained task space and in the generalization task space (Fig 7: gray bars vs. post-training white bars). Moreover, we observed no clear difference in *testing* block performances between the trained task space and the generalization task space. By contrast, comparison of the *cont/exp_xy_* index between the baseline 1 (pre-training) and baseline 2 (post-training) blocks on Day 1 revealed a training effect i.e., a significant reduction toward the ideal value for the outer targets [T_14_ = 5.432; p < 0.0001] and the inner targets [T_14_=3.9876; p = 0.0013] (Fig 7; compare white bars on either side of the vertical dotted line for Day 1). Taken together, these results support the conclusion that practicing to use supplemental limb state feedback improved subjects’ short-term internal representations of extrapersonal space throughout the arm’s workspace, even when that feedback was no longer present.

**Figure 7:**
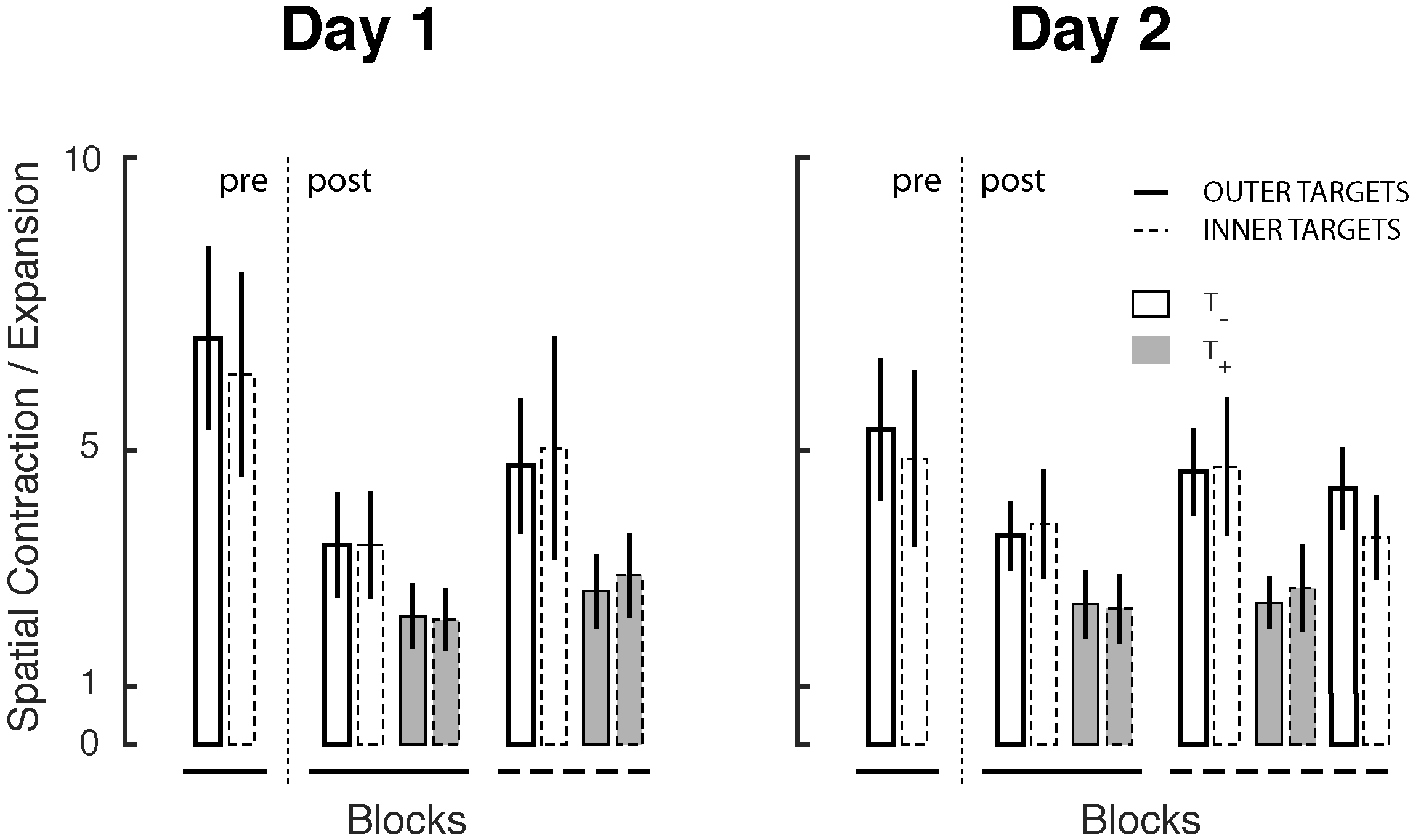
Group results for spatial contraction/expansion vs. blocks in experimental phases wherein visual cues were precluded (V_−_). Pre-training: *baseline 1*; post-training: *baseline 2* and *testing* in the trained workspace (white and gray bars above solid horizontal line), *baseline 3* and *testing* in the generalization workspace (white and gray bars above dashed horizontal line). Error bars: 95% CI; solid bars: *count/exp_xy_* computed for the outer targets; dashed bars: *count/exp_xy_* computed for the inner targets; Gray bars: blocks with vibrotactile feedback (T_+_); White bars: blocks without vibrotactile feedback (T_−_).

### Aspect Ratio

We next scrutinized hand path aspect ratio to evaluate the extent to which training with vibrotactile feedback facilitated the production of rectilinear hand paths. Hand paths were relatively straight when subjects were provided visual feedback of hand motion (Fig 8, black bars; aspect ratios less than 0.2). On the contrary, paths exhibited marked curvature in the presence of supplemental vibrotactile feedback (Fig 8, gray bars; with aspect ratios greater than 0.4). Hand path curvatures assumed intermediate values with neither visual nor vibrotactile feedback. The fact that aspect ratio increased when vibrotactile feedback was provided suggests that subjects employed different control strategies to reach the intended target depending on current feedback conditions (we explore this observation further, below). Moreover, we observed no meaningful difference between aspect ratios produced in the initial and final training blocks on either day, suggesting that subjects may need much more than two training sessions if they are to recover “natural” straight-line hand trajectories when incorporating vibrotactile limb state feedback into the on-going control of reaching movements.

**Figure 8:**
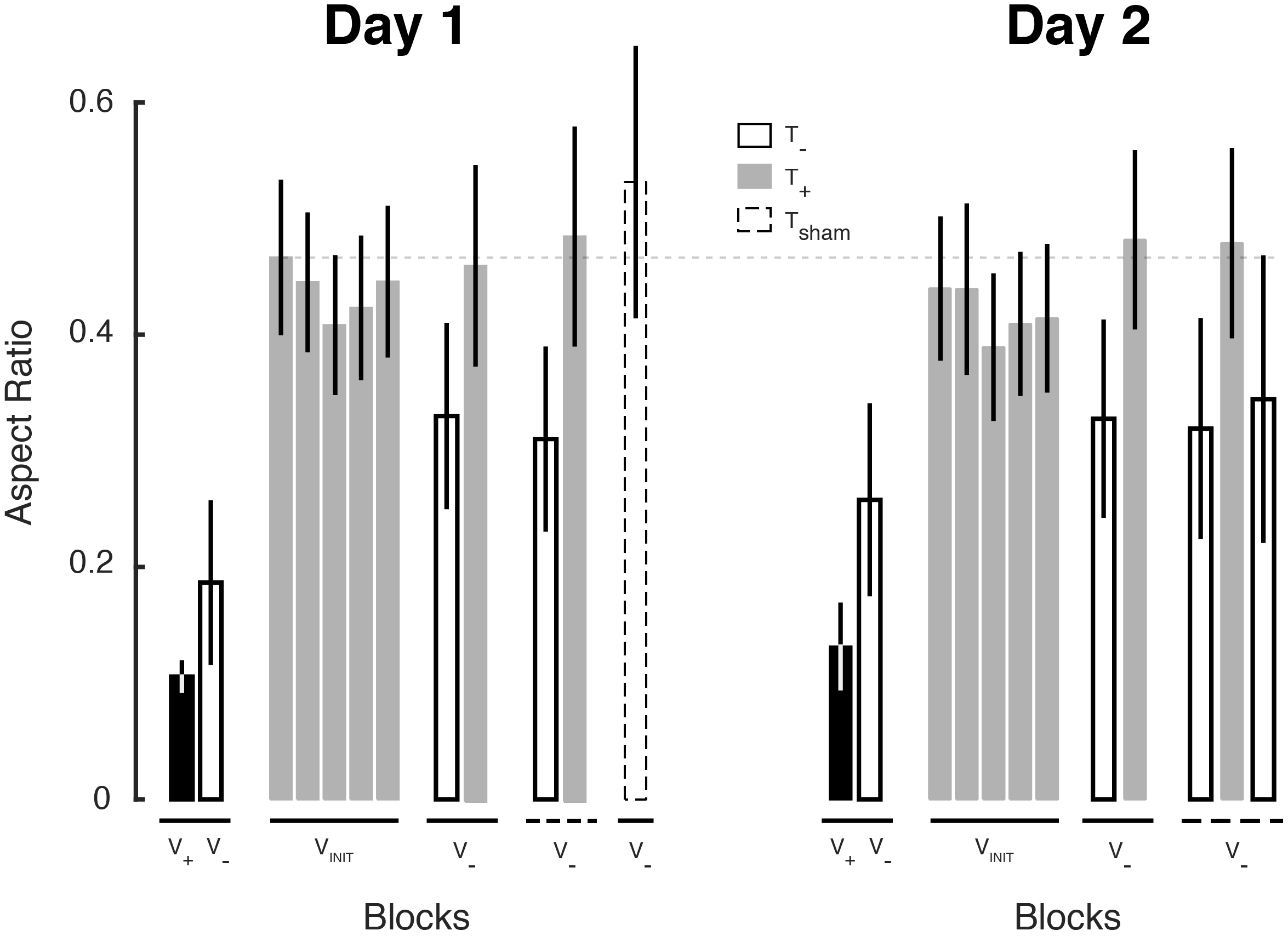
Group results for aspect ratio vs. blocks for both days. Error bars: 95% CI; Black bars: blocks with visual feedback (V_+_). V_−_ indicates blocks without visual feedback. Gray bars: blocks with vibrotactile feedback (T_+_); White bars: blocks without vibrotactile feedback (T_−_); dashed bar: blocks with sham feedback (T_sham_).

### Decomposition Index

Kinematic analyses of individual subject hand trajectories during each experimental phase reveal the use of different movement strategies prior to training with the vibrotactile feedback vs. after training. Prior to training, subjects made goal-directed reaches that were reasonably straight (Fig 9A, a representative baseline movement shown with dashed line), with hand speed profiles that were approximately bell-shaped (Fig 9B top; thin dashed trace). By contrast, movements performed after training typically had bimodal hand speed profiles (Fig 9B bottom; thin solid trace) corresponding to hand paths that first moved predominantly along one cardinal axis of the vibrotactile display, then the other. The selected baseline and training hand paths shown in Figure 9A had unitless DI values (Eqn 1) equal to 0.40 and 0.82, respectively.

**Figure 9:**
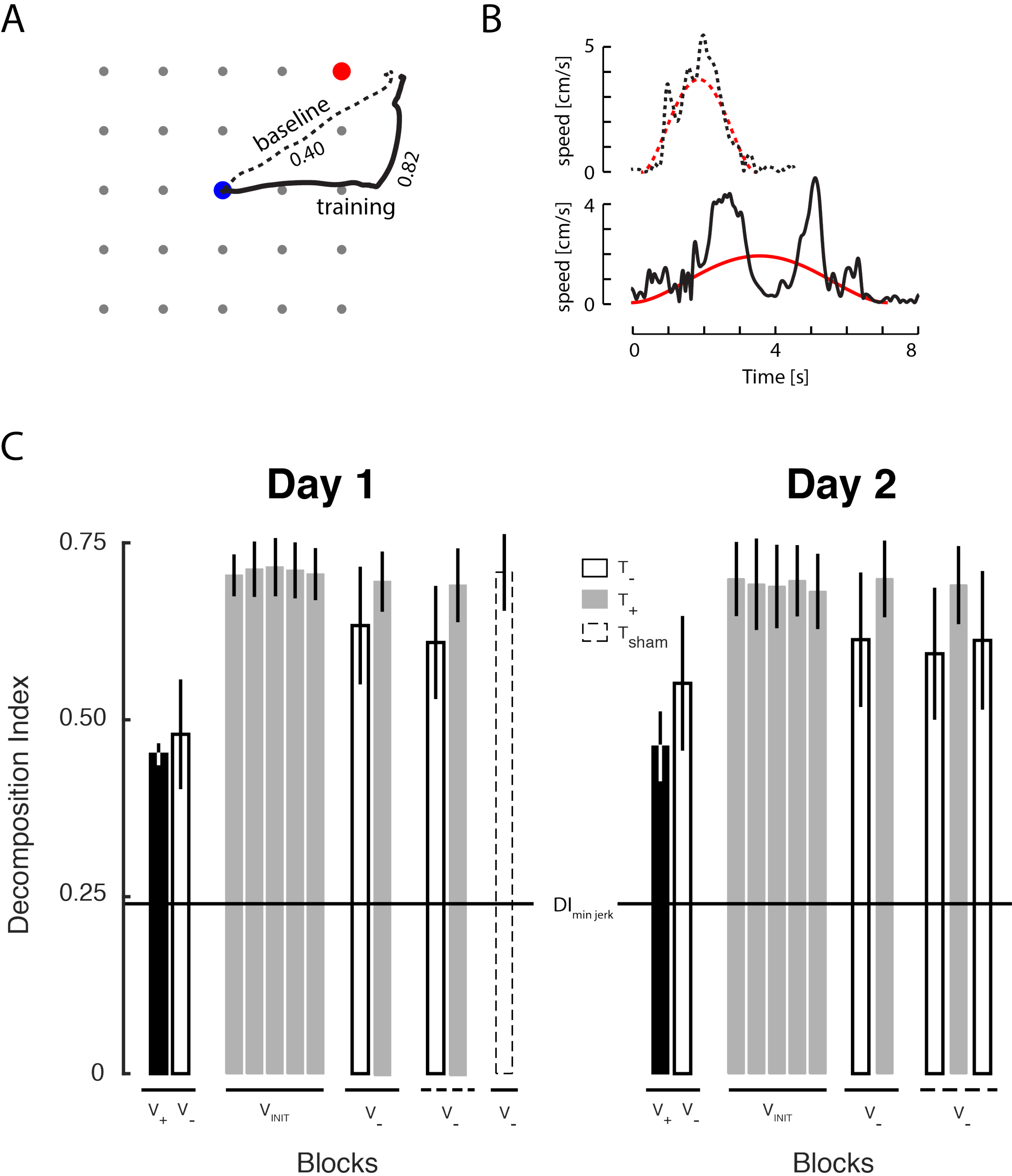
Decomposition Index (DI) vs. experimental block. A) Sample hand paths, and B) the corresponding speed profiles performed by a selected subject from a movement during the baseline 1 block (dashed traces) and the first training block (solid traces). On initial exposure to the reach task, the subject made movements that were reasonably straight (A) with speed profiles that were roughly bell-shaped (B). Also plotted for comparison in (B) are speed profiles from ideal minimum-jerk trajectories (thick black traces). Whereas the minimum-jerk trajectories always yield DI values equal to 0.24, regardless of movement extent, movement speed, and movement direction (excepting the singular cases described in Appendix 1), the baseline block movement with single speed peak yielded a DI value of 0.40 whereas the training block movement with two speed peaks yielded a DI value of 0.82. C) Group results for DI values vs. experimental block for both days. The meanings of bar shadings and labels are consistent with those described for Figure 8. The long horizontal reference bar corresponds to the DI value obtained from ideal straight-line minimum-jerk movements. Error bars: 95% CI.

As described in the APPENDIX, off-axis minimum-jerk hand paths yield DI values equal to 0.24 (Fig 9C; horizontal line). This is true regardless of movement extent, movement speed, or movement direction (excepting trajectories directed along one of the cardinal axes of the vibrotactile display, which yield singular DI values; see the APPENDIX for more details). Numerical DI values for baseline movements deviate from the ideal value because individual baseline movements had hand trajectories that were not quite straight and hand speed profiles that were not quite smooth. By contrast, DI values for training trajectories were relatively high because hand paths were curved and speed profiles were bimodal.

The kinematic observations presented for individual trajectories (Figs 9A and 9B) generally held true across the study population across both days of training (Fig 9C). Whereas DI values were relatively low prior to training with vibrotactile feedback on Day 1, DI values immediately jumped high at the onset of training as subjects employed a “decomposed” movement strategy whereby they first moved along one cardinal axis of the vibrotactile display, then the other. At the onset of training, all 15 subjects made vibrotactile-guided reaches with DI values greater than or equal to 0.60. By the end of Day 2 training, 13 of 15 subjects persisted in making vibrotactile-guided reaches with DI values greater than 0.60. We did not observe any meaningful reduction in DI values across the two days of training (i.e., from the first training block on Day 1 to the last training block on Day 2; T_14_ = 0.9039, p = 0.381). Remarkably, decomposition persisted in post-training reaches performed without vibrotactile feedback, both in the trained and contralateral workspaces on both days

### Target Capture Time

Subjects were instructed to capture the targets as accurately as possible, without constraint on time or hand path. We expected that initial attempts to integrate the novel vibrotactile feedback signals into the ongoing planning and control of arm movement would increase target capture time as subjects initially learned how best to interpret and use the vibrotactile information, but that target capture times would decrease as learning progressed. We therefore examined target capture time across the different feedback conditions to verify that subjects did in fact attempt to integrate the vibrotactile feedback into their ongoing planning and control rather than to ignore it. We found a marked increase in target capture time of about 2 seconds when supplemental sensory feedback was provided (Fig 10, gray bars), suggesting that the integration of vibrotactile feedback into the ongoing planning and control of limb motion involved considerable cognitive processing relative to movements performed without that feedback. A comparison of initial training performance across days reveals that target capture time was longer on Day 1 relative to Day 2 (T_14_ = 2.6205, p < 0.05); although the magnitude of this decrease was quite small, the results add additional support to the conclusion that subjects did retain on Day 2 some aspects of what they learned while training to use supplemental sensory feedback on Day 1.

**Figure 10:**
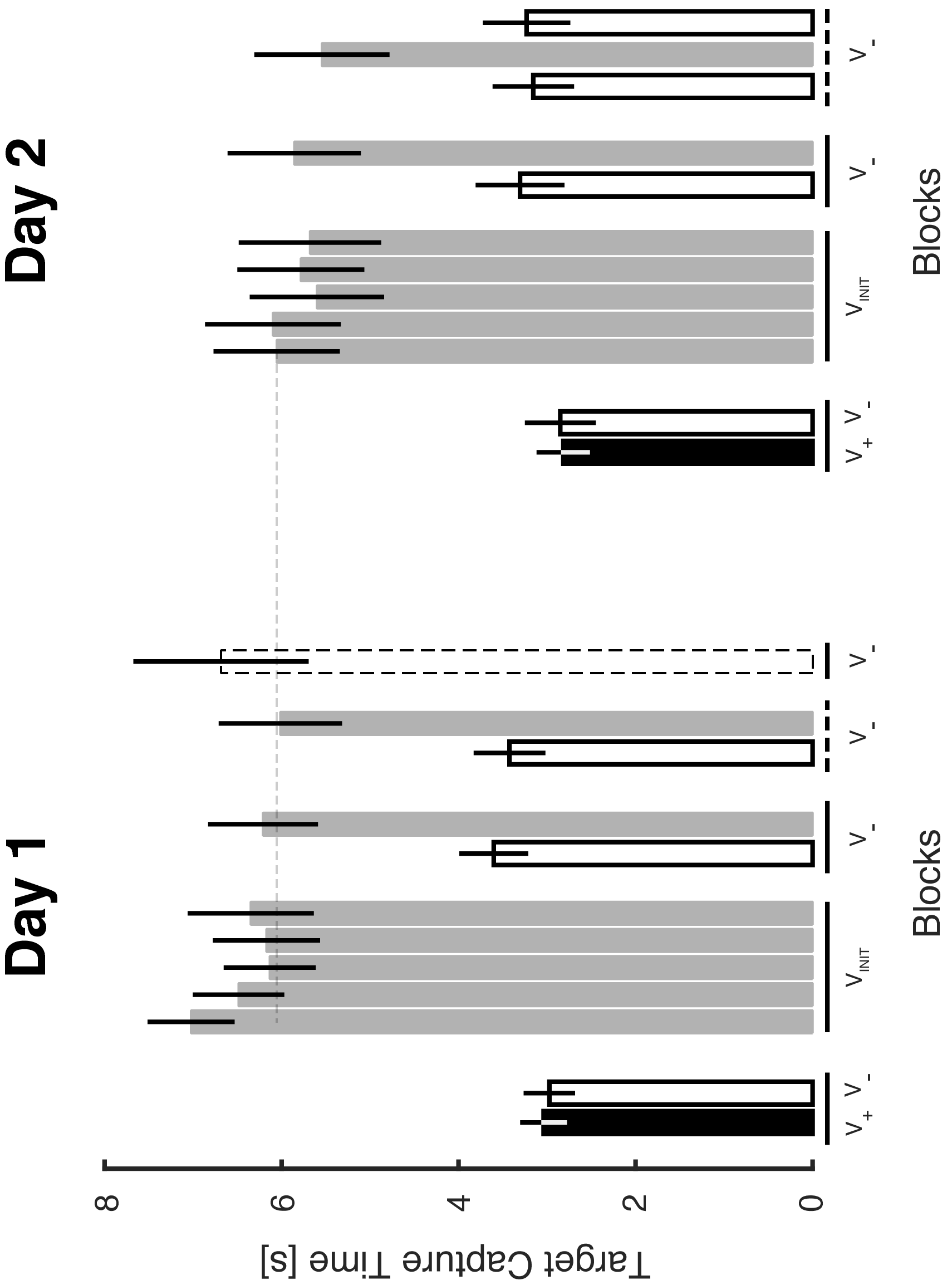
Group results for target capture time vs blocks for both days. Error bars: 95% CI; Black bars: blocks with visual feedback (V_+_). V_−_ indicates blocks without visual feedback. Gray bars: blocks with vibrotactile feedback (T_+_); White bars: blocks without vibrotactile feedback (T_−_); dashed bar: sham feedback (T_sham_). Horizontal dashed line: visual guideline for comparing initial training blocks across days.

We observed no difference in target capture time between the sham condition and other blocks with vibration. This outcome suggests that although sham-block performance degraded dramatically relative to trials when informative vibration was provided, subjects did in fact attempt to use the sham feedback to drive performance and did not simply ignore it.

## Discussion

This study tested the application of vibrotactile stimulation as supplemental state feedback for enhancing the on-going control of a moving limb. Subjects wore a two-dimensional vibrotactile display on their non-dominant arm while performing horizontal planar reaching movements with their dominant arm. We constructed a mapping between the position of the moving hand and its representation within the vibrotactile display such that small, goal-directed hand displacements were more easily discriminable using supplemental vibrotactile feedback than with intrinsic proprioceptive feedback in the moving arm (cf. Shah et al. 2016; Fuentes and Bastian, 2010). After mere minutes of training, subjects were able to use the information encoded within the vibrotactile feedback to perform blind reaches (i.e. to capture the visual target without concurrent cursor feedback of hand position). Subjects performed with levels of accuracy and precision that exceeded levels attained without the supplemental feedback (i.e., using intrinsic proprioceptive feedback alone). Reach accuracy continued to improve within and across two days of training with additional practice on using supplemental vibrotactile feedback. These improvements were due to partial resolution of distortions in the internal representation of reachable space (i.e., proprioceptive drift and task space expansion) that spontaneously arose when concurrent cursor feedback of hand motion was eliminated. We also observed a beneficial aftereffect of vibrotactile feedback training (a reduction of task space expansion) when the vibrotactile feedback was turned off; this observation was consistent with the hypothesis that supplemental vibrotactile limb state feedback training can improve internal representations of extrapersonal space. Finally, the utility of vibrotactile feedback was not limited to the trained workspace; subjects immediately capitalized on the information encoded within the vibrotactile feedback to perform blind reaches in an unpracticed region of the arm’s workspace.

In both workspaces, all subjects immediately adopted a movement strategy by which they attempted to capture targets by moving first along one cardinal axis of the vibrotactile display and then along the other (i.e., a “decomposition” strategy). Kinematic consequences of decomposition included elevated levels of hand path curvature and bimodal hand speed profiles after training. The decomposition strategy was persistent; 13 of 15 subjects continued to decompose their movements after two days of training. Thus, this limited amount of training did not suffice to allow subjects to fully integrate supplemental limb state feedback into the planning and control of goal-directed reaching such that the movements have straight hand paths and unimodal hand speed profiles (cf. Morasso, 1981). The fact that post-training reaches reflected the decomposition strategy even when vibrotactile feedback was turned off indicates that at least part of the persistence was due to altered movement plans, and not just to a change in the way the brain uses concurrent sensory information to guide the ongoing feedback control of movement. Future studies are warranted to identify training programs that will encourage integration of vibrotactile limb state feedback into the planning and control of goal-directed movements with straight hand paths and unimodal hand speed profiles. The results of such studies will have practical implications for the augmentation of human performance in a variety of applications where precision and efficiency of manual movements is desired, including rehabilitation of functional movement after stroke and the enhancement of skilled manual performance in teleoperation tasks such as robotic surgery.

### Vibratory feedback of limb state enhances the online control of movement beyond limits imposed by intrinsic proprioception

Subjects could reach to each of the visual targets with accuracy and precision when cursor feedback of hand position was provided, but dramatic drift of reach endpoints toward the midline and expansion of task space along both planar dimensions of task space occurred rapidly after cursor feedback was extinguished in the first baseline block of trials. These reach errors reflect distortions of an internal representation of reachable space (i.e., task space and/or body image) that evolve within a time frame of seconds. Remarkably, all subjects were able to reduce these distortions after mere minutes of prior exposure to the vibrotactile display and the information it encoded (see Fig 5; day 1, training block 1). To do so, subjects had to integrate a completely novel set of limb-state-dependent stimuli into their plans for movement and its ongoing control. Although target capture errors decreased further both within and across two days of training, target capture times did not decrease meaningfully over the two days, suggesting that this limited training was insufficient for subjects to transition far beyond the initial cognitive stage of sensorimotor skill learning and into the associative phase (c.f., Fitts and Posner 1967). These two phases rely on the so-called “body image”, an internal representation of the body that is accessible to consciousness (see Proske and Gandevia, 2012, for a description and discussion of the concept of body image). Further studies should explore how extended and structured training schedules can impact the rate at which integration of vibrotactile limb state feedback into the ongoing control of goal-directed movements progresses through the later, “automatic” phase of sensorimotor skill learning. The later phases of skill learning involve a separate internal representation of the body (the body schema) that works independently of consciousness (c.f., Proske and Gandevia, 2012).

Two novel aspects of our experimental approach likely contributed to the rapid learning observed in our study (i.e., the reduction of drift and the reduction in the expansion of endpoints seen Figs 4, 6, and 7). First and foremost, vibrotactile feedback encoded limb state information in our study rather than performance error information, as used by most previous studies of sensory augmentation via vibrotactile stimulation (e.g., Bark et al., 2015; Cuppone et al., 2016). Whereas limb state encoding preserves a bijective (one-to-one) relationship between hand position in the workspace and the stimulus presented by the vibrotactile display, error encoding does not. Limb state encoding maintains consistency in the relationships between visual, proprioceptive, and the novel vibrotactile signals, such that initial movement trials could establish cross-modal spatial links between unimodal representations of initial or desired hand positions (c.f., Deneve and Pouget, 2004), and later trials could reinforce those relationships. By contrast, we would not expect that encodings of task performance error would induce persistent changes in internal representation(s) of reachable space because the relationships between visual, proprioceptive, and vibrotactile signals would reset each time the initial starting conditions and/or desired goal targets change. Second, the lateral displacement of the physical workspaces from the visual workspace probably accentuated the evolution of hand position drift toward the midline in the absence of both visual and vibrotactile feedback.

In any case, beneficial effects of vibrotactile limb state feedback reflect its partial integration into the online control of movement because some benefits were seen only while the vibrotactile signals were present; upon removal of the vibrotactile signals on Day 1, target capture errors increased, as did the amount of drift and the area spanned by the reach endpoints. The ability to integrate vibrotactile limb state feedback into ongoing control of movement was not limited to the trained workspace because the effects of vibration were reported also in the untrained contralateral workspace. When subjects were provided with concurrent vibrotactile limb state feedback on Day 2, target capture errors again decreased, as did directional and extent errors as well as did the phenomenon of spatial expansion. Because performances degraded during the last generalization baseline in the untrained workspace on Day 2, we reject the possibility that the learning effect experienced with concurrent vibration feedback was due merely to more prolonged practice on the reaching task.

Rapid changes in the internal representation of reachable space have also been observed in experimental studies using single unit neural recordings to explore tool use, and in studies of crosslesional extinction in neurological populations. In one exemplar study, Iriki and colleagues trained macaque monkeys to retrieve distant objects using a rake, while they recorded the activity of neurons in the caudal postcentral gyrus, where somatosensory and visual signals converge (Iriki et al., 1996). The activities of these cells, and of others in premotor cortex (see Graziano and Gross, 1995) code for tactile events on a body part-such as the hand-as well as for visual events near that body part. Iriki and colleagues found that the visual receptive fields of bimodal neurons were altered during tool use to cover the expanded accessible space, potentially representing neural correlates of a modified body schema in which the tool was incorporated into an internal representation of the hand (Iriki et al., 1996). Other studies have exploited a deficit known as crosslesional extinction in patients with right hemisphere lesions of the frontal and parietal cortices to show that tool-use elicits analogous expansion of the space within which the human brain represents reachable visual targets (see Farne and Ladavas, 2000; Maravita et al., 2001; also see di Pellegrino and Ladavas 2015 for a recent review). Taken together, such studies suggest that the brain employs malleable central representations of *peripersonal space* – the “reachable” region of space that is in close proximity to the body, and that the convergence of multimodal sensory information onto single cells within association and premotor regions of cortex contributes importantly to those central representations. We speculate that convergence of multimodal sensory information (vibrotactile and visual) onto single cells within association and premotor regions of cortex may underlie the rapid integration of novel vibrotactile limb state signals into the planning and ongoing control of movement.

### Vibrotactile feedback training elicits adaptive changes in motor plans

Vibrotactile limb state feedback training had aftereffects that resulted not only in short-term improvements in the internal representation of extrapersonal space, but in lasting changes in motor plans for reaching. When subjects were not provided visual or supplemental feedback about their movements, they made large movements that yielded large target capture errors and an expansion of the workspace spanned by the hand at the end of the reaches. After training, in the same reduced feedback conditions (i.e., with only proprioceptive feedback available), subjects exhibited improved reach accuracy and reduced spatial expansion indicating that the training had had a persistent and beneficial impact on the sensorimotor transformation(s) relating visual targets to desired reach endpoints. The finding that subjects also persisted in decomposing their movements along the cardinal axis of the vibrotactile display in the post-training baseline blocks demonstrates that subjects applied to these movements the same movement strategy adopted when the vibrational feedback was provided.

Cuppone and colleagues have also found a persistent improvement of motor performance after a training based on concurrent vibrotactile and haptic feedback (Cuppone et al., 2016). In that study, groups of subjects grasped the handle of wrist manipulandum and were tested pre-and post-training in their ability to perform a 2-DOF (flexion/extension, abduction/adduction) position matching task and a 2-DOF target tracking task. One group received small forces (haptic cues) that guided the hand to the target during training. Two additional groups received the haptic cues along with 1-DOF vibrotactile cues related to the “lateral deviation” of the wrist from the straight line connecting a neutral home position to the desired 2-DOF target. The two groups differed according to the delivery site of vibrotactile stimulation. A control group received no training. Relative to the untrained control group, proprioceptive acuity was improved after training only in the groups that received both the haptic and vibrotactile cues. Improvement did not depend on which limb received the vibrotactile cues. The groups that received vibrotactile cues exhibited improved movement kinematics during goal-directed wrist movements from the onset of training (i.e., reduced lateral deviations), although improvement came at the cost of increased movement times and an increased number of “movement units” (i.e., velocity profile peaks) required to capture the target. Since the haptic and vibrotactile cues were not tested in isolation but presented concurrently, Cuppone et al. could not determine whether haptic force cues or vibratory feedback of movement errors were necessary elements of the proprioceptive training they described.

The results of the current study support and extend the findings of Cuppone and colleagues (Cuppone et al., 2016). Also in our case, observed improvements in accuracy and representation of the workspace could be due both to the vibrational feedback and to the terminal haptic cues that we used to provide knowledge of results during training (i.e., robotic translation of the hand to the intended final target positions). However, in the present study, the motor plan adopted by the subjects after training changed profoundly, presenting clear carryover effects of movement strategies adopted while subjects attended to the vibrational feedback. We believe the decomposition strategy was motivated primarily by the vibrotactile limb state feedback for four reasons. First, the decomposition strategy arose during the part of the movement that was not impacted by the terminal haptic feedback. Second, submovements of the decomposition strategy were aligned primarily along the cardinal axes of the vibrotactile display. Third, terminal haptic feedback was provided only after subjects had used the vibrotactile feedback to correct for movement errors and had indicated that they had captured the target. Finally, the terminal haptic feedback corrected for target capture errors by driving the subject’s hand straight to the target, not along a decomposed path.

The finding that subjects used a stereotyped, strategic approach toward integrating vibrotactile feedback into the planning and execution of goal-directed reaches is interesting *per se*. When performing goal-directed reaches, the hand typically follows a relatively straight path between its initial and final positions, with a unimodal hand speed profile that is often described as “bell-shaped” (Morasso 1981; Abend et al., 1982; Flash and Hogan 1985; Sergio and Scott, 1998). Such point-to-point trajectories are planned to be straight in visually perceived space (Wolpert et al., 1995), and can become markedly curved either when a nonlinear transformation is interposed between the motion of the end effector (hand) and its visual representation (Flanagan and Rao, 1995) or when accounting for the geometrical properties of the object that the brain is controlling (Danziger and Mussa-Ivaldi, 2012). In our case, all 15 subjects appeared to employ the same, unusual decomposition strategy to capture targets in absence of visual feedback and in presence of informative vibrotactile feedback of hand position. Instead of treating vibrotactile feedback of hand position as a vector quantity, i.e., by simultaneously reducing target capture error along both axes of the vibrotactile display, subjects first reduced error along one dimension and then along the other. Most subjects self-reported at the conclusion of the experiments that they had attempted to “process one feedback dimension at a time” or that they had planned hand movements “so that only one [pair of tactors] would provide meaningful vibration at any given time.” Furthermore, two subjects explicitly reported that the subjacent gray grid underneath the targets was a useful cue to interpret the vibrotactile feedback, suggesting that they were able to “feel” their way along the grid to the desired target. The grid we display might have influenced the adoption and retention of the decomposition strategy, but it could not have been the main contributing factor since that strategy emerged only when vibrational feedback was provided. Moreover, movement decomposition was not merely a reflection of how the vibrotactile feedback was processed in real-time, but also must have reflected, in part, a training-dependent alteration in the movement plans, because decomposition persisted in post-training reaches in both the trained and contralateral workspaces-whether or not supplemental vibrotactile feedback of hand position was currently available. Future studies should: explore the factors motivating adoption and retention of a decomposition strategy in training with multi-DOF vibrotactile displays; identify training strategies to encourage the vectorial interpretation of the vibrotactile stimuli and the simultaneous reduction of target capture error along all axes of the vibrotactile display (e.g., the production of goal-directed movements with straight hand paths and unimodal hand speed profiles); and assess the utility of vibrotactile sensory substitution systems and appropriate training strategies to promote successful completion of typical daily tasks by suitable patient populations, such as those survivors of stroke who have deficits of proprioceptive sensation and yet retain residual movement capability.

## Appendix

Using the Matlab script presented in the listing below, we simulated ideal, “minimum-jerk” hand trajectory (Flash & Hogan, 1985) in the horizontal plane. As expected for a dimensionless performance measure, we found the value of the Decomposition Index of Equation 1 to be invariant across movement direction, movement extent, and movement speed for these straight-line hand movements. Figure 11A depicts the trajectories produced by one ‘run’ of the script with N = 16 movement directions, a movement extent of 10 cm, and a movement duration of 5 seconds (Fig 11B, solid trace). Each of the simulated reaches yielded a DI value of 0.24. We also simulated movements having durations of 3 seconds (Fig 11B, dashed trace); these also yielded DI values of 0.24. Identical values were found for movements in all directions when we increased or decreased movement length, or varied the number of movements simulated (data not shown). The only exception to these general observations were trajectories for which there was no motion along one or the other of the cardinal directions, in which case the Decomposition Index of Equation 1 becomes singular. Because real-world hand trajectories will in all likelihood have some motion along both the X and Y axes, the practical utility of the DI for evaluating human performance is unconstrained by the exceptional cases.

**Figure 11 (Appendix):**
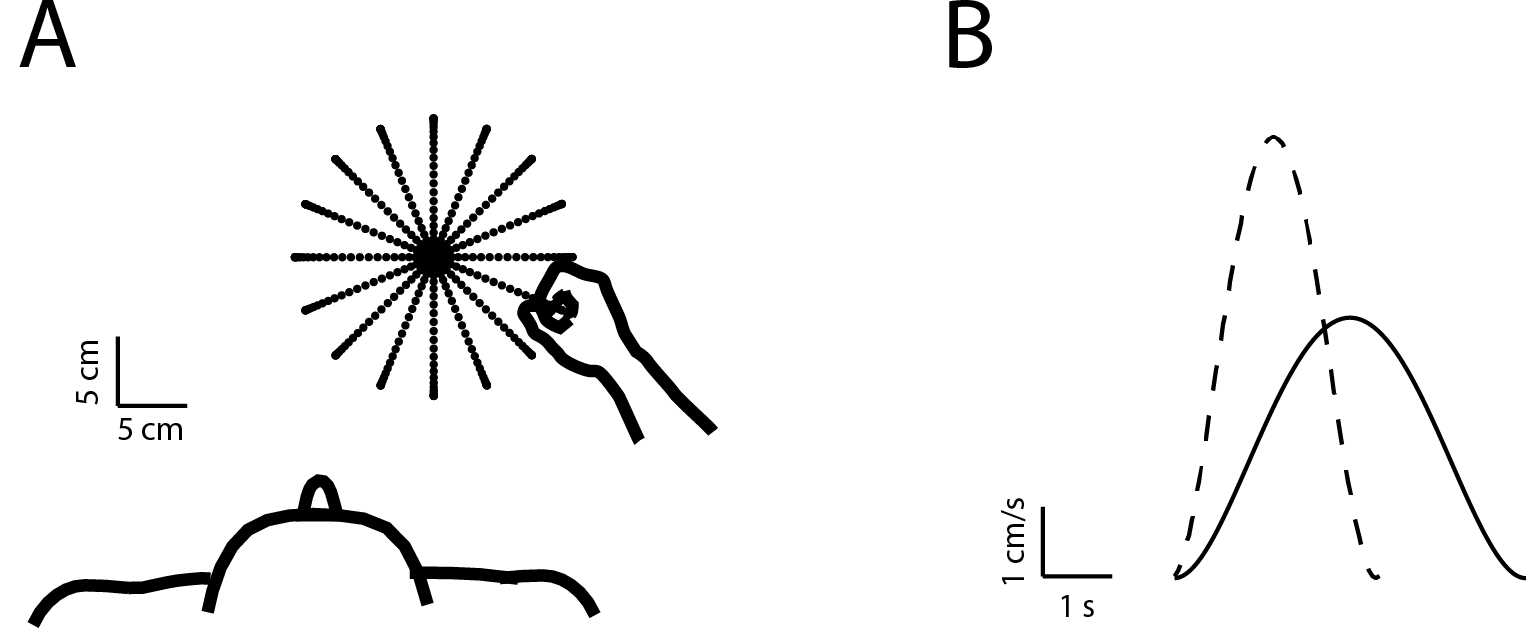
Simulation analysis of the Decomposition Index (DI) of Equation 1. A) Each of the sixteen 10 cm minimum-jerk hand trajectories yielded (unitless) DI values of 0.238. B) Identical DI values were obtained from movements of different speeds, and different extents. Here, two simulated hand speed profiles are presented, corresponding to reaches lasting 3 seconds and 5 seconds. Identical DI values are obtained for much faster reaches (data not shown).

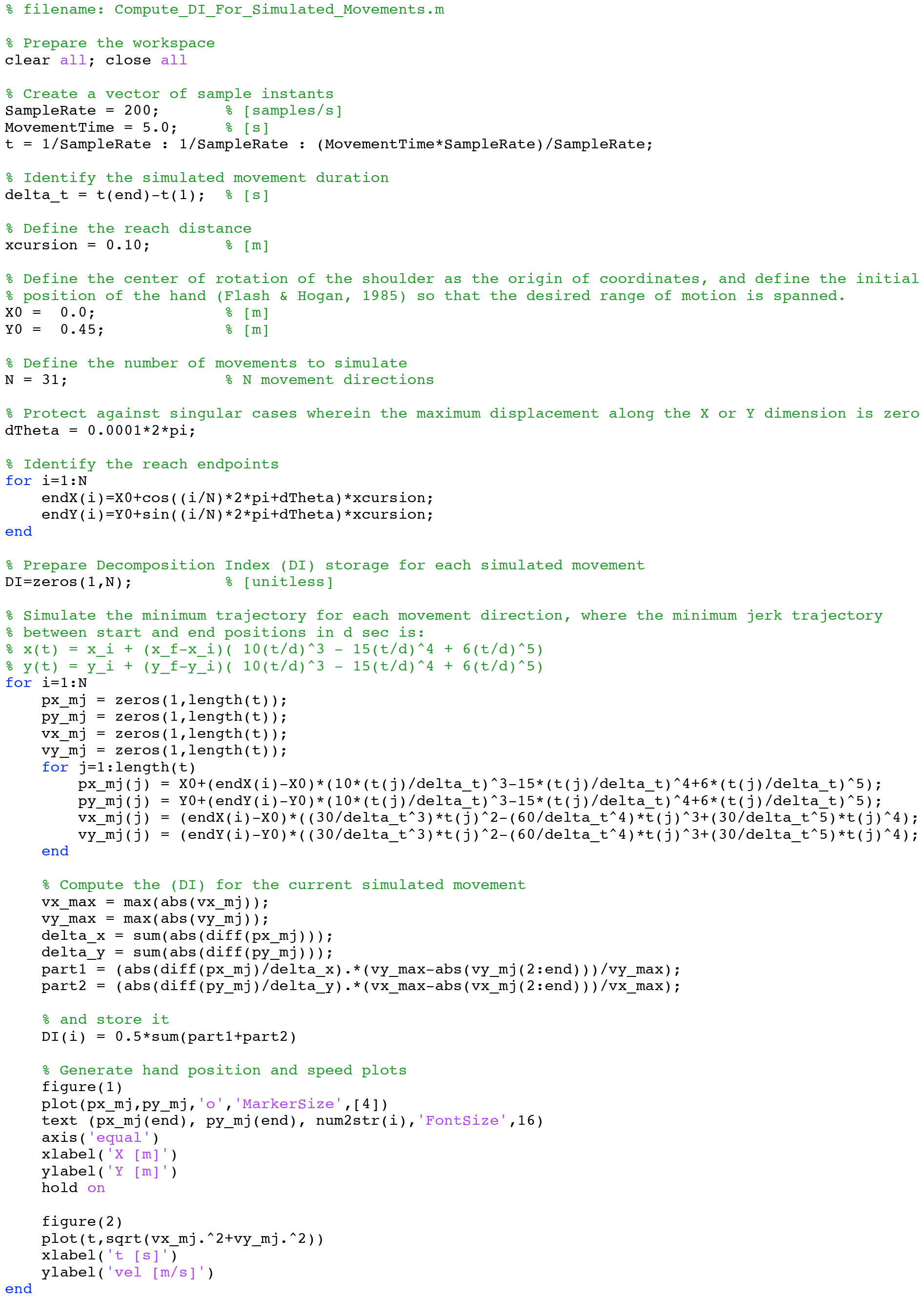

**Code Listing 1:** Matlab script for simulating minimum-jerk hand trajectories and computing resulting Decomposition Index (DI) values.

## References

Abend, W., Bizzi, E., & Morasso, P. (1982). Human arm trajectory formation. Brain: a journal of neurology, 105(Pt 2), 331–348.

Bark, K., Khanna, P., Irwin, R., Kapur, P., Jax, S. A., Buxbaum, L. J., & Kuchenbecker, K. J. (2011). Lessons in using vibrotactile feedback to guide fast arm motions. Paper presented at the World Haptics Conference (WHC), 2011 IEEE.

Bark, K., Hyman, E., Tan, F., Cha, E., Jax, S. A., Buxbaum, L. J., & Kuchenbecker, K. J. (2015). Effects of vibrotactile feedback on human learning of arm motions. IEEE Transactions on Neural Systems and Rehabilitation Engineering, 23(1), 51–63.

Bastian, H. C. (1887). On Different Kinds of Aphasia, with Special Reference to Their Classification and Ultimate Pathology. Br Med J, 2(1401), 985–990.

Blennerhassett, J. M., Matyas, T. A., & Carey, L. M. (2007). Impaired discrimination of surface friction contributes to pinch grip deficit after stroke. Neurorehabil Neural Repair, 21(3), 263–272. doi:10.1177/1545968306295560

Cameron, B. D., de la Malla, C., & Lopez-Moliner, J. (2014). The role of differential delays in integrating transient visual and proprioceptive information. Front Psychol, 5, 50. doi:10.3389/fpsyg.2014.00050

Carey, L. M., & Matyas, T. A. (2011). Frequency of discriminative sensory loss in the hand after stroke in a rehabilitation setting. J Rehabil Med, 43(3), 257–263. doi:10.2340/16501977-0662

Connell, L. A., Lincoln, N. B., & Radford, K. A. (2008). Somatosensory impairment after stroke: frequency of different deficits and their recovery. Clin Rehabil, 22(8), 758–767. doi:10.1177/0269215508090674

Cuppone, A., Squeri, V., Semprini, M., & Konczak, J. (2015). Robot-assisted training to improve proprioception does benefit from added vibro-tactile feedback. Conf Proc IEEE Eng Med Biol Soc, 2015, 258–261. doi:10.1109/EMBC.2015.7318349

Cuppone, A. V., Squeri, V., Semprini, M., Masia, L., & Konczak, J. (2016). Robot-assisted proprioceptive training with added vibro-tactile feedback enhances somatosensory and motor performance. PloS one, 11(10), e0164511.

Danziger, Z., & Mussa-Ivaldi, F. A. (2012). The influence of visual motion on motor learning. Journal of Neuroscience, 32(29), 9859–9869.

di Pellegrino, G., & Làdavas, E. (2015). Peripersonal space in the brain. Neuropsychologia, 66, 126–133.

De Santis, D., Zenzeri, J., Casadio, M., Masia, L., Riva, A., Morasso, P., & Squeri, V. (2015). Robot-assisted training of the kinesthetic sense: enhancing proprioception after stroke. Frontiers in human neuroscience, 8, 1037.

Flash, T., & Hogan, N. (1985). The coordination of arm movements: an experimentally confirmed mathematical model. J Neurosci, 5(7), 1688–1703.

Dukelow, S. P., Herter, T. M., Moore, K. D., Demers, M. J., Glasgow, J. I., Bagg, S. D., … Scott, S. H. (2010). Quantitative assessment of limb position sense following stroke. Neurorehabil Neural Repair, 24(2), 178–187. doi:10.1177/1545968309345267

Farnè, A., & Làdavas, E. (2000). Dynamic size-change of hand peripersonal space following tool use. Neuroreport, 11(8), 1645–1649.

Fitts, P. M., & Posner, M. I. (1967). Human performance.

Flanagan, J. R., & Rao, A. K. (1995). Trajectory adaptation to a nonlinear visuomotor transformation: evidence of motion planning in visually perceived space. Journal of neurophysiology, 74(5), 2174–2178.

Flash, T., & Hogan, N. (1985). The coordination of arm movements: an experimentally confirmed mathematical model. Journal of neuroscience, 5(7), 1688–1703.

Fuentes, C. T., & Bastian, A. J. (2010). Where is your arm? Variations in proprioception across space and tasks. J Neurophysiol, 103(1), 164–171. doi:10.1152/jn.00494.2009

Gandevia, S. C., McCloskey, D. I., & Burke, D. (1992). Kinaesthetic signals and muscle contraction. Trends Neurosci, 15(2), 62–65.

Ghez, C., Gordon, J., & Ghilardi, M. F. (1995). Impairments of reaching movements in patients without proprioception. II. Effects of visual information on accuracy. J Neurophysiol, 73(1), 361–372. doi:10.1152/jn.1995.73.1.361

Gordon, J. A. M. E. S., Ghilardi, M. F., & Ghez, C. (1995). Impairments of reaching movements in patients without proprioception. I. Spatial errors. Journal of neurophysiology, 73(1), 347–360.

Graziano, M. S., & Gross, C. G. (1995). The representation of extrapersonal space: a possible role for bimodal, visual-tactile neurons. The cognitive neurosciences, 1021–1034.

Iriki, A., Tanaka, M., & Iwamura, Y. (1996). Coding of modified body schema during tool use by macaque postcentral neurones. Neuroreport, 7(14), 2325–2330.

Krueger, A. R., Giannoni, P., Shah, V., Casadio, M., & Scheidt, R. A. (2017). Supplemental vibrotactile feedback control of stabilization and reaching actions of the arm using limb state and position error encodings. J Neuroeng Rehabil, 14(1), 36. doi:10.1186/s12984-017-0248-8

Lee, M. W. L., McPhee, R. W., & Stringer, M. D. (2008). An evidence-based approach to human dermatomes. Clinical Anatomy, 21(5), 363–373.

Lieberman, J., & Breazeal, C. (2007). TIKL: Development of a wearable vibrotactile feedback suit for improved human motor learning. IEEE Transactions on Robotics, 23(5), 919–926.

Maravita, A., Husain, M., Clarke, K., & Driver, J. (2001). Reaching with a tool extends visual–tactile interactions into far space: Evidence from cross-modal extinction. Neuropsychologia, 39(6), 580–585.

Matthews, P. B. (1988). Proprioceptors and their contribution to somatosensory mapping; complex messages require complex processing. Canadian journal of physiology and pharmacology, 66(4), 430–438.

Morasso, P. (1981). Spatial Control of Arm Movements. Experimental Brain Research, 42,223–227.

Paillard, J., & Brouchon, M. (1968). Active and passive movements in the calibration of position sense. The neuropsychology of spatially oriented behavior, 11, 37–55.

Proske, U., & Gandevia, S. C. (2012). The proprioceptive senses: their roles in signaling body shape, body position and movement, and muscle force. Physiol Rev, 92(4), 1651–1697. doi:10.1152/physrev.00048.2011

Risi N, Krueger A, Giannoni P, Casadio M, Scheidt RA (2016) Learning to use supplemental kinaesthetic feedback for enhancing reach performance. Karniel Computational Motor Control Workshop, Beer-Sheva, Israel.

Risi N, Mrotek LA, Shah V, Casadio M, Scheidt RA (2017) Learning to use supplemental vibrotactile feedback of limb position enhances goal-directed reach performance. Soc. Neural Control Movement, Dublin, Ireland.

Sainburg, R. L., Poizner, H., & Ghez, C. (1993). Loss of proprioception produces deficits in interjoint coordination. J Neurophysiol, 70(5), 2136–2147. doi:10.1152/jn.1993.70.5.2136

Sanes, J. N., Mauritz, K. H., Evarts, E. V., Dalakas, M. C., & Chu, A. (1984). Motor deficits in patients with large-fiber sensory neuropathy. Proc Natl Acad Sci U S A, 81(3), 979–982.

Sarlegna, F. R., Gauthier, G. M., Bourdin, C., Vercher, J. L., & Blouin, J. (2006). Internally driven control of reaching movements: a study on a proprioceptively deafferented subject. Brain Res Bull, 69(4), 404–415. doi:10.1016/j.brainresbull.2006.02.005

Scheidt, R. A., Conditt, M. A., Secco, E. L., & Mussa-Ivaldi, F. A. (2005). Interaction of visual and proprioceptive feedback during adaptation of human reaching movements. J Neurophysiol, 93(6), 3200–3213. doi:10.1152/jn.00947.2004

Scheidt, R. A., & Stoeckmann, T. (2007). Reach adaptation and final position control amid environmental uncertainty after stroke. J Neurophysiol, 97(4), 2824–2836. doi:10.1152/jn.00870.2006

Scheidt, R. A., Lillis, K. P., & Emerson, S. J. (2010). Visual, motor and attentional influences on proprioceptive contributions to perception of hand path rectilinearity during reaching. Experimental brain research, 204(2), 239–254.

Sergio, L. E., & Scott, S. H. (1998). Hand and joint paths during reaching movements with and without vision. Experimental brain research, 122(2), 157–164.

Shah, V., Gagas, M., Krueger, A., Iandolo, R., Peters, D., Casadio, M., & Scheidt, R. (2016). Vibrotactile discrimination thresholds vary among dermatomes in the upper extremity of healthy humans. San Diego: Society for Neuroscience.

Smeets, J. B., van den Dobbelsteen, J. J., de Grave, D. D., van Beers, R. J., & Brenner, E. (2006). Sensory integration does not lead to sensory calibration. Proc Natl Acad Sci U S A, 103(49), 18781–18786. doi:10.1073/pnas.0607687103

Sober, S. J., & Sabes, P. N. (2003). Multisensory integration during motor planning. J Neurosci, 23(18), 6982–6992.

Tyson, S. F., Hanley, M., Chillala, J., Selley, A. B., & Tallis, R. C. (2008). Sensory loss in hospital-admitted people with stroke: characteristics, associated factors, and relationship with function. Neurorehabil Neural Repair, 22(2), 166–172. doi:10.1177/1545968307305523

Tzorakoleftherakis, E., Bengtson, M. C., Mussa-Ivaldi, F. A., Scheidt, R. A., & Murphey, T. D. (2015). Tactile proprioceptive input in robotic rehabilitation after stroke. Paper presented at the Robotics and Automation (ICRA), 2015 IEEE International Conference on.

Wann, J. P., & Ibrahim, S. F. (1992). Does limb proprioception drift? Exp Brain Res, 91(1), 162–166.

Wolpert, D. M., Ghahramani, Z., & Jordan, M. I. (1995). Are arm trajectories planned in kinematic or dynamic coordinates? An adaptation study. Experimental brain research, 103(3), 460–470.

Wong, J. D., Wilson, E. T., & Gribble, P. L. (2011). Spatially selective enhancement of proprioceptive acuity following motor learning. J Neurophysiol, 105(5), 2512–2521. doi:10.1152/jn.00949.2010

Zackowski, K. M., Dromerick, A. W., Sahrmann, S. A., Thach, W. T., & Bastian, A. J. (2004). How do strength, sensation, spasticity and joint individuation relate to the reaching deficits of people with chronic hemiparesis? Brain, 127(Pt 5), 1035–1046. doi:10.1093/brain/awh116

